# ARBUR, a machine learning-based analysis system for relating behaviors and ultrasonic vocalizations of rats

**DOI:** 10.1101/2023.12.19.572288

**Authors:** Zhe Chen, Guanglu Jia, Qijie Zhou, Yulai Zhang, Zhenzhen Quan, Xuechao Chen, Toshio Fukuda, Qiang Huang, Qing Shi

## Abstract

Deciphering how different types of behavior and ultrasonic vocalizations (USVs) of rats interact can yield insights into the neural basis of social interaction. However, the behavior-vocalization interplay of rats remains elusive because of the challenges of relating the two communication media in complex social contexts. Here, we propose a machine learning-based analysis system (ARBUR) that can cluster without bias both non-step (continuous) and step USVs, hierarchically detect eight types of behavior of two freely behaving rats with high accuracy, and locate the vocal rat in 3-D space. By simultaneously recording the video and ultrasonic streams of two freely behaving rats, ARBUR reveals that rats communicate via distinct USVs when engaging in different types of social behavior. Moreover, we show that ARBUR can not only automatically reveal the well-understood behavior-associated vocalizations that were carefully concluded by other behavioral researchers, but also hold the promise to indicate novel findings that can be hardly found by manual analysis, especially regarding step USVs and the active/passive rat-associated USVs during easy-to-confuse social behaviors. This work highlights the potential of machine learning algorithms in automatic animal behavioral and acoustic analysis and could help mechanistically understand the interactive influence between the behaviors and USVs of rats.

## Introduction

Nonverbal communication and vocalization are two vital means of natural communication during social interaction across the animal kingdom and human society ***Barker et al.*** (***2021***); ***Padilla-Coreano et al.*** (***2022***). Nonverbal communication such as facial expression ***Dolensek et al.*** (***2020***), body posture, waggle dance ***Dong et al.*** (***2023***), and social play ***VanRyzin et al.*** (***2019***) can be received by nearby conspecifics and often analyzed generally by human observers as social behavior ***Robinson et al.*** (***2008***), whereas vocalization can be communicated remotely to convey information in dark and tortuous environments. Both social behavior and vocalization can be readily recorded experimentally as outer observable states that reflect the inner emotional states ***Dolensek et al.*** (***2020***) or even neural circuit activities ***Wei et al.*** (***2021***); ***Li et al.*** (***2022b***); ***Schneider et al.*** (***2023***), and therefore are actively investigated in animal behavior research ***Murugan et al.*** (***2017***). During social engagement, both the social behavior and vocalization of individuals influence each other in a continuous and interactive manner, contributing to the variable group social dynamics. Deciphering the behavior-vocalization interplay ***Knutson et al.*** (***1998***); ***Sangiamo et al.*** (***2020***) would thus reveal insights into the neural basis of social interaction. For such deciphering efforts, rats are widely used as social animal models because of their innate sociability ***Venniro and Shaham*** (***2020***) and proven emotion-related ultrasonic vocalizations (USVs) ***Brudzynski*** (***2013***). For example, the alarm sounds (22-kHz USVs) of trapped rats may induce overwhelming distress in freely moving rats via emotional contagion and further evoke pro-social behavior in the free ones ***Bartal et al.*** (***2011***). However, a dedicated system for analyzing social communications of freely behaving rats is still missing to promote the mechanistic understanding of the interplay between social behavior and USVs.

The barrier is caused by the dfficulty of relating the different types of behavior and USVs of rats in complex social contexts (to reveal the underlying behavior-vocalization interplay) because of the following challenges. First of all, a method for the unbiased and automatic clustering of USVs that not only covers both non-step (continuous) and step (non-continuous) signals in spectrograms ***Sangiamo et al.*** (***2020***) but also comprehensively incorporates the structure of frequency and duration ***Takahashi et al.*** (***2010***), is still missing. Moreover, the automatic behavior detection of rats under social interaction poses a major challenge in constructing the behavior-specific features ***Lorbach et al.*** (***2018***); ***Vogt*** (***2021***); therefore, it is difficult to discriminate different types of easy-to-confuse social behavior ***Bohnslav et al.*** (***2021***); ***Harris et al.*** (***2023***); ***Segalin et al.*** (***2021***); ***Kabra et al.*** (***2013***). In addition, allocating the recorded USVs to the vocal rat also brings an obstacle because of the proximity and top-view partial overlapping of socially engaged rats ***Neunuebel et al.*** (***2015***).

In this Article, we propose a machine learning-based Analysis system for Relating Behaviors and USVs of Rats (named ARBUR). ARBUR automatically clusters ultrasonic syllables into user-defined or automatically calculated subgroups from a comprehensive perspective, detects eight types of behavior (Table 1) based on the lateral camera view, and locates the vocal rat in two-rat scenarios with free social interaction. Using ARBUR, we reveal that the distinct behaviors of rats are associated with different USVs. Therefore, based on the simultaneously recorded raw video and audio streams (see Figure 2–figure supplement 1), ARBUR can relate the behavior and USVs of rats in freely interacting social contexts for downstream qualitative and quantitative analyses. For example, we show that the pinned rat produces significantly more 22-kHz aversive USVs compared with the bully one or during other social behaviors or solitary states. Moreover, ARBUR indicates several novel findings about still-associated or moving-associated USVs, which have long been neglected in the manual analysis-dominated research.

**Table 1.** Behavioral definitions of freely behaving rats.

| Behavior | Definition |
| --- | --- |
| SO (solitary) | Both rats engage in separate activities. |
| ST (still) | Both rats keep still. |
| MA (moving away) | A rat moves away from another rat. |
| AP (approaching) | A rat moves towards another rat until contact. |
| FO (following) | A rat follows another rat without contact. |
| PIN (pinning) | A rat holds down another rat lying on its back. |
| POU (pouncing) | A rat lies on the back of another rat. |
| SNC (social nose contact) | A rat touches another rat with its nose tip. |

## Results

### Unsupervised clustering of the vocal repertoire

We recorded 43,357 ultrasonic vocalizations (USVs) to create the vocal repertoire. To cluster the USVs in an unbiased and comprehensive way, we present the unsupervised three-step clustering algorithm ARBUR:USV. In the first step, ARBUR:USV divides the USVs into 2 clusters according to their mean frequency-indicated emotional states (aversive 22-kHz USVs (ranging from 15 kHz to 32 kHz) or appetitive 50-kHz USVs (above 32 kHz)). This is a special treatment for rat USVs analysis since rats produce meaningful USVs with a frequency near 22 kHz while mice don’t. Another special consideration in ARBUR:USV is the inclusion of step signals, which are usually overlooked in other research. Moreover, note that 50-kHz USVs are classified into four groups according to their mean frequency and duration: high-peak-frequency (HPF) 50-kHz USVs and low-peak-frequency (LPF) 50-kHz USVs, because two aggregated areas were observed in the density map (Figure 2b), indicating two kinds of 50-kHz USVs may be present despite of their similar frequency contours (shape in the spectrograms). Therefore, in the second step, the 50-kHz USVs are classified into four groups (groups A-D) according to their continuity (non-step or step) and distribution in frequency-duration space (HPF or LPF) (Figure 2a). In the third step, ARBUR:USV separates the four 50-kHz groups (groups A-D) into subgroups within each group according to their frequency contours, thus allowing users to investigate the correlation between USV contours with other biological factors (e.g. behavior types) in relevant research. In the meantime, the 22-kHz group (group E) is divided into five subgroups based on mean frequency and duration because the frequency contours of 22-kHz USVs do not vary much.

While the first two steps divide the USVs according to their biological relevance (mean frequency, natural distribution in frequency-duration space), the third clustering step separates the USVs within each group according to their contour features, which raises the question of determining the optimal number of clusters to balance over-clustering with under-clustering. Here, ARBUR:USV provides three solutions for users to choose. First, the number of clusters for groups A-E can be arbitrarily chosen according to their experiences. This may be suitable for experienced animal behaviorist to investigate USV-related rat behaviors with fixed cluster size. Second, ARBUR:USV can calculate the optimal number of clusters for each group according to the elbow method (see Methods) ***Coffey et al.*** (***2019***). It basically relies on the relationship (curve) of total within-cluster error (TWCE) versus the increasing number of clusters. With increasing cluster numbers, TWCE tends to decrease exponentially. The elbow method finds the elbow point of the curve to be the optimal number. For the collected dataset, the optimal numbers for groups A-E are 13, 28, 13, 14, and 8, respectively (see Figure 2–figure supplement 2). Third, ARBUR:USV also provides the progressive method, which considers the clustering degree for each cluster (see Methods). In particular, the cluster number, starting with two, increases by one if the average innerpoint percent of clustering results does not reach a user-defined threshold. For example, if the threshold is set to be 0.98, the optimal cluster number for group E is 8 (Figure 2–figure supplement 3). It can produce a higher or lower number of clusters if stricter or looser thresholds are used for particular purposes. In short, ARBUR:USV provides three methods for choosing the number of clusters for groups A-E. Here, to balance simpler visualization and the elbow method result, we arbitrarily set the numbers for groups A-E to be 10, 25, 10, 15, and 5, respectively.

Figure 2a shows the clustering examples. Note that the USV clusters were re-sorted in descending order according to contour slope (for clusters 1-60) or duration (for clusters 61-65). It is intuitive that the USVs within each cluster share similar contours, and the mean contours of each cluster differ from those of the others (Figure 2–figure supplement 4). We then quantified the clustering results. We show that ARBUR:USV achieves much higher inter-cluster distance compared with intra-cluster variance across the 65 clusters (Figure 2c; Figure 2–figure supplement 5a, b, c). Moreover, we quantified the 65 clusters of the USVs, and show that they vary substantially in terms of duration, frequency range, mean frequency, and signal counts across clusters (Figure 2–figure supplement 6).

### Hierarchical behavior classification of two freely behaving rats

ARBUR:Behavior is based on a hierarchical classification architecture, which we optimized for analyzing the species-specific behaviors of two freely behaving rats (see Methods). We constructed different classification features based on behavioral states (non-social, moving, and social) and designed three classifiers to distinguish behavioral categories further (see Figure 3–figure supplement 1). ARBUR:Behavior uses the binocular side-view video stream (segmented by SegFormer ***Xie et al.*** (***2021***) to remove the background) as input to discriminate eight behaviors of rats (Figure 1b). ARBUR:Behavior can also run in a single-shot mode if moving behaviors are not considered.

**Figure 1.**
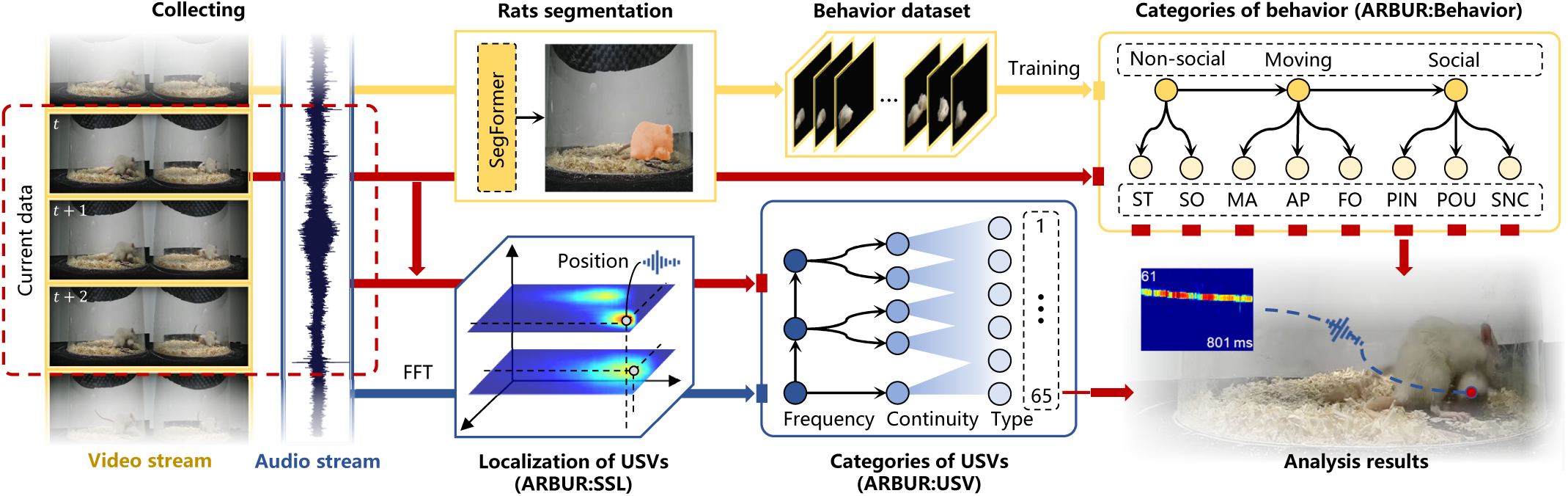
Illustration of the working flow of ARBUR. ARBUR takes simultaneously recorded video and audio streams as input and outputs the current behavior type, the ultrasonic vocalization (USV) in spectrogram with cluster type and duration indicated, and the labeled position of the vocal rat for each frame of the binocular images in the video frame. **Figure 1—figure supplement 1.** Experimental setup for recording video and audio streams of rats simultaneously.

**Figure 2.**
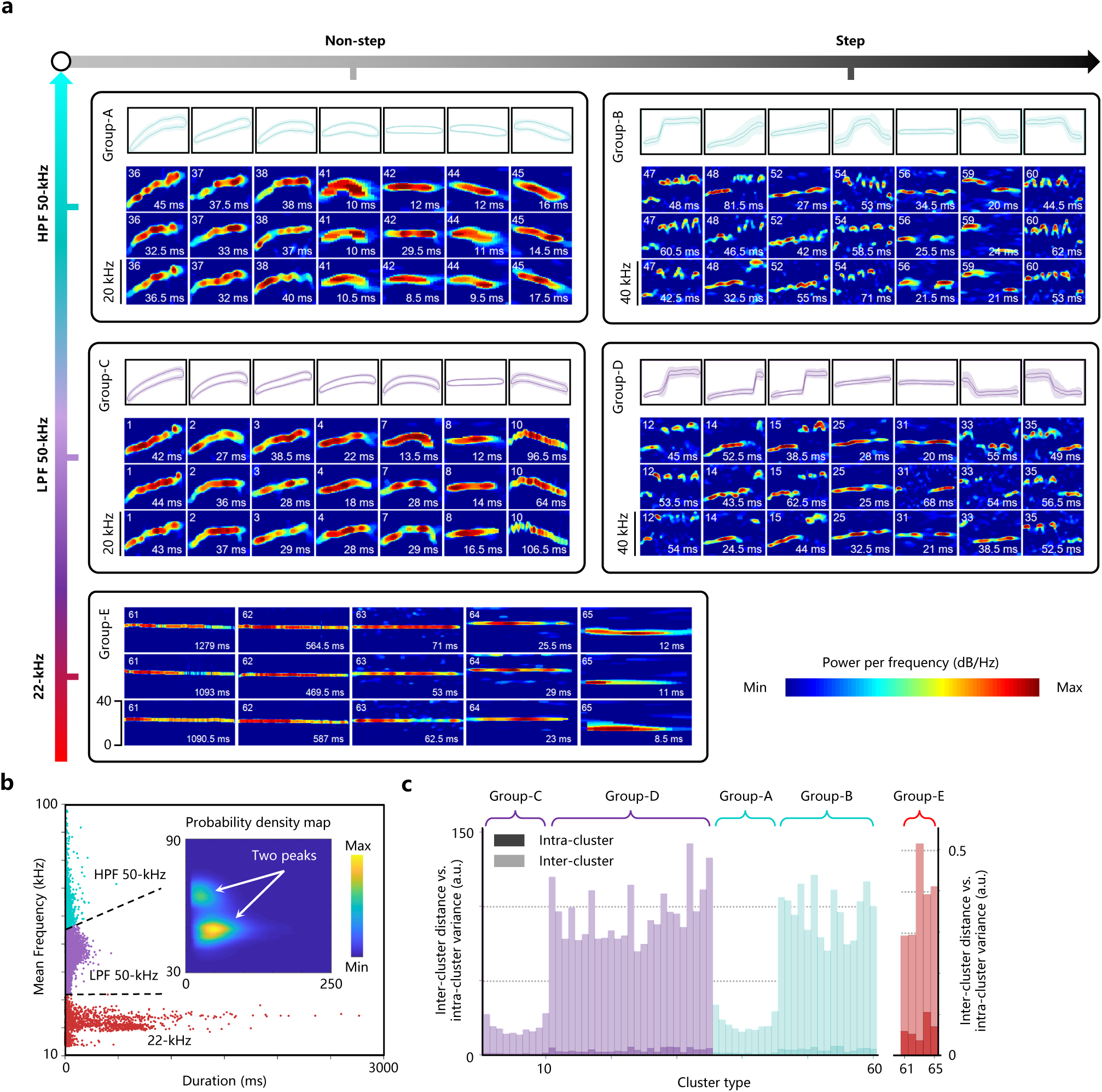
ARBUR:USV clusters the vocal repertoire in an unbiased and comprehensive manner. **a**, Representative examples show that USVs within the same clusters share similar contours. Groups A-D, top panels, mean value of contour ± s.e.m. Bottom three panels, three selected USVs spectrograms with cluster type (top left) and duration (bottom right) are indicated. Each column represents one cluster. Intensity is indicated by color (red, maximum; blue, minimum). Groups A-E represent high-frequency non-step 50-kHz USVs (clusters 36-45), high-frequency step 50-kHz USVs (clusters 46-60), low-frequency non-step 50-kHz USVs (clusters 1-10), low-frequency step 50-kHz USVs (clusters 11-35) and 22-kHz USVs (clusters 61-65), respectively. The frequency range of spectrograms: 20 kHz for group A and C; 40 kHz for group B and D; 0-40 kHz for group E. **b**, Intuitive demonstration of the classification within the 50-kHz USVs, made because two independent peaks were observed in the probability density map (inset). **c**, Quantification of the clustering results, which show high inter-cluster distances and small intra-cluster variances. **Figure 2—figure supplement 1.** Workflow of the unsupervised clustering algorithm ARBUR:USV. **Figure 2—figure supplement 2.** Determining the optimal number of clusters for groups A-E using the elbow method in our dataset. **Figure 2—figure supplement 3.** Determining the optimal number of clusters using the progressive method for group E. **Figure 2—figure supplement 4.** Mean value (solid lines) ± s.e.m. (shadow) of frequency contours within each cluster for clusters 1-60. **Figure 2—figure supplement 5.** Validating ARBUR:USV. **Figure 2—figure supplement 6.** Quantification of the clustering results.

We annotated 1,265 randomly extracted segments from all recorded videos (including 43,357 segments) to test the performance of ARBUR:Behavior. As demonstrated in Figure 3a, the module can classify the behaviors of two freely behaving rats well, especially the moving behaviors with large displacement and the social behaviors with complex postures. We show that the desired detection precision, recall, and F1-score of each behavior are achieved (average precision: 0.874, average recall: 0.862, average F1-score: 0.868, Figure 3b). The confusion matrix shows that ARBUR:Behavior can discriminate the easy-to-confuse social behaviors with high F1-scores, such as PIN (pinning) and POU (pouncing) (Figure 3c). In contrast to studies based on end-to-end architectures ***Bohnslav et al.*** (***2021***); ***Marks et al.*** (***2022***); ***Gerós et al.*** (***2022***), ARBUR:Behavior is capable of discriminating specific social behaviors (PIN, POU, and SNC (social nose contact) and can classify a higher number of behavior categories while achieving comparable accuracy.

**Figure 3.**
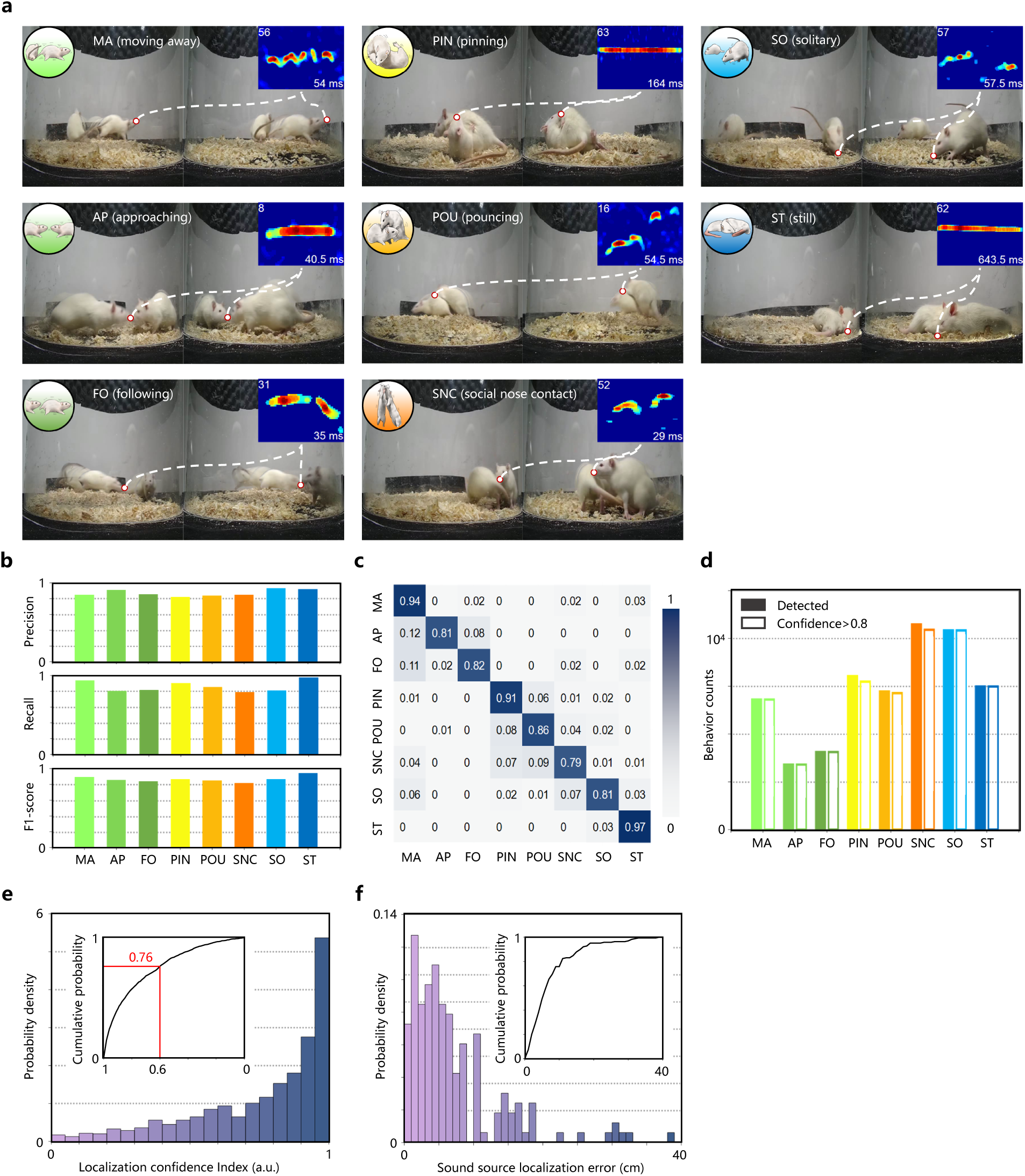
ARBUR simultaneously achieves accurate behavior detection and sound source localization in three dimensions. **a**, Examples of the ARBUR outputs for eight behaviors: left and right camera view, behavior type (top left of each panel), spectrograms of USVs with cluster (inset, top left), and duration (inset, bottom right) and location of the vocal rat in the two camera views (red circle). Frequency range of spectrograms: 20 kHz for cluster 8; 0-40 kHz for clusters 62 and 63; 40 kHz for all other clusters. **b**, Quantification of the performance of ARBUR:Behavior. **c**, Confusion matrix showing the identification rates of ARBUR:Behavior. **d**, Counts of different types of behavior detected (logarithmic) across behavior types. **e**, Distribution of the localization confidence index (LCI) obtained by ARBUR:SSL; inset, cumulative probability vs. LCI. **f**, Distribution of the sound source localization error by ARBUR:SSL; inset, cumulative probability vs. error. **Figure 3—figure supplement 1.** Workflow of ARBUR:Behavior. **Figure 3—figure supplement 2.** Comparison of algorithms for discriminating the easy-to-confuse social state behaviors. **Figure 3—figure supplement 3.** Workflow of ARBUR:SSL. **Figure 3—figure supplement 4.** Quantitative evaluation of the snout-snout distance distribution for rat-rat free interaction scenarios. **Figure 3—figure supplement 5.** Validating ARBUR:SSL.

In addition, we tested the performance of current mainstream deep learning classification algorithms ***Touvron et al.*** (***2021***); ***Liu et al.*** (***2021***, 2022); ***He et al.*** (***2016***) for the direct classification of the eight behaviors on our dataset. ResNet achieves the best performance with an accuracy of only 0.333 using an eight-class classifier (Figure 3-figure supplement 2a). For the two- and three-class classification of three social behaviors, the highest accuracies of these algorithms are 0.767 and 0.578 (Figure 3-figure supplement 2b). We argue that features with behavioral specificity can not be directly extracted in the images using the end-to-end approach, resulting in poor classification performance. Therefore, some studies extracted the pose and tracking data of rats to classify behaviors in an unsupervised way and achieved high accuracies ***Li et al.*** (***2022a***); ***Hsu and Yttri*** (***2021***). However, such methods are unable to discriminate specific social behaviors. In contrast, ARBUR:Behavior detects non-social behaviors using the location feature of rats in the image and moving behaviors by estimating the optical flow of the rat as well as its relative orientation. To discriminate social behaviors, it extracts “histogram of oriented gradients” (HOG) ***Dalal and Triggs*** (***2005***) descriptors to represent the statistics of social posture features in a standardized way.

We also tested the performance of mainstream machine learning-based classifiers for two- and three-class classifiers on three social behaviors (Figure 3–figure supplement 2b). The support vector machine (SVM) achieved the highest classification accuracy (0.843) in binary classification, and the Receiver Operating Characteristic (ROC) curve shows that it has the best overall classification performance (Figure 3-figure supplement 2c). Therefore, we designed an SVM-based decision tree classifier for discriminating social behaviors (see Methods). ARBUR:Behavior has a greater advantage in terms of F1-score/accuracy and the number of behavior categories, compared with hand-crafted features used in existing studies ***Harris et al.*** (***2023***); ***Jhuang et al.*** (***2010***); ***Segalin et al.*** (***2021***). On balance, ARBUR:Behavior outperforms existing studies with respect to comprehensive detection performance (Table 3). ARBUR:Behavior successfully detected 33,428 behaviors (with a confidence of over 0.8) out of 43,357 segments of video frames corresponding to recorded USVs (Figure 3d).

### Locating the vocal rat in 3-D space

To locate the vocal rat during free-behaving scenarios, we developed a novel algorithm (ARBUR:SSL) that incorporates the lateral binocular view (for rat nose reconstruction in the Cartesian space), behavior classification results, height-sensitive distributed microphone configuration, and 3-D probabilistic triangulation of sound source to locate the vocal rat in 3-D space (see Figure 3–figure supplement 3). ARBUR:SSL takes the 3D reconstructed nose position, the current behavior type, and four-channel audio streams as input. It can comprehensively leverage these information and output the pixel position of the vocal rat in both binocular images, which is useful for follow-up expert quantitative and qualitative analysis.

The rat-rat social interaction scenario makes the microphone array far above the rats (50 mm above ground) compared with the experimental sets for mouse interaction, which makes the recorded USVs signal intensity (therefore signal-to-noise ratio) much weaker than others. This poses a new challenge to us because the mainstream sound source localization (SSL) algorithms suitable for mouse interaction experiments work much worse in our dataset and, therefore undesirable for our purposes. To address this challenge, we optimized a classic method ***Neunuebel et al.*** (***2015***) and proposed a new index (localization confidence index, LCI), which measures the consistency between 3-mic SSL and 4-mic SSL results, and hence can rule out those low-SNR USVs. Using the proposed method, we achieved a median SSL error of 5.01 cm in the test dataset, which is suitable for rat-rat social scenarios. Since the large size of adult rats (>18 cm), snout-to-snout distance in rat-rat interaction scenarios is much bigger than in mouse-mouse interactions (Figure 3–figure supplement 4). Such an SSL error can be usable for safely assigning the USVs to the vocal rat in rat-rat interaction scenarios.

Another concern that haunted the experts working on SSL is the lack of accurate test datasets. Maintream studies reportedly induced mouse USVs using heterosexual urine ***Sterling et al.*** (***2023***); ***Oliveira-Stahl et al.*** (***2023***), but the visual localization of rodent snout would bring unnecessary errors, and lack the ability of snout height detection. In this paper, we constructed a rat USV test dataset for evaluating SSL errors using USVs collected from still rats, which brings two advantages: 1) all the USVs are produced at the same spatial source position because the rat kept still throughout the process; 2) 3D position of the vocal rat nose/snout is provided, therefore allowing evaluating 3D SSL algorithms.

We further show that a higher LCI is associated with smaller sound source localization (SSL) error (Figure 3–figure supplement 5b). Therefore, only USV signals with an LCI over 0.6 are accepted, which consist of 76% of the valid vocal repertoire (29772 USVs, intersection of the valid outputs of ARBUR:USV and ARBUR:Behavior, Figure 4–figure supplement 1). ARBUR:SSL shows decent horizontal-plane localization precision, with less than 10 cm error for over 78% of the data, which is considerable, considering the much weaker USVs in our dataset, compared to existing algorithms (see ***Neunuebel et al.*** (***2015***); ***Sangiamo et al.*** (***2020***)) (Figure 3f, Figure 3–figure supplement 5a), and more importantly, suitable for our purposes. Moreover, by evaluating the LCI at different heights (reconstructed from the binocular view), ARBUR:SSL can pinpoint the vocal rat even if the two rats overlap in the top-down projection plane (which is more frequent in, for example, pinning and pouncing). ARBUR:SSL assigned 17,235 USVs out of 29,772 USVs to the vocal rat. Figure 3a shows examples. In contrast to other sound source localization methods for rodents’ USVs, ARBUR:SSL features the ability to locate the vocal rat in 3-D space while preserving the localization precision with weaker USV signals (Table 4).

### Revealing the behavior-vocalization interplay

By seamlessly incorporating these three modules, ARBUR can build connections between different types of rat behavior and vocalizations. Figure 4a shows examples (also see Movie S1). It is intuitive that when the rats are engaged in non-aggressive play behavior, such as social nose contact and pouncing, more appetitive 50-kHz USVs are communicated. On the contrary, the submissive one emits more aversive 22-kHz USVs when engaged in aggressive play behavior, i.e., pinning. We investigated whether these two hypotheses would hold in the whole repertoire. Therefore, we quantified the USV proportions of rats in different types of behavior (Figure 4b). Results show that the submissive rat produces significantly more aversive 22-kHz USVs during the pinning process. Moreover, in social nose contact and pouncing, both rats produce more 50-kHz USVs.

**Figure 4.**
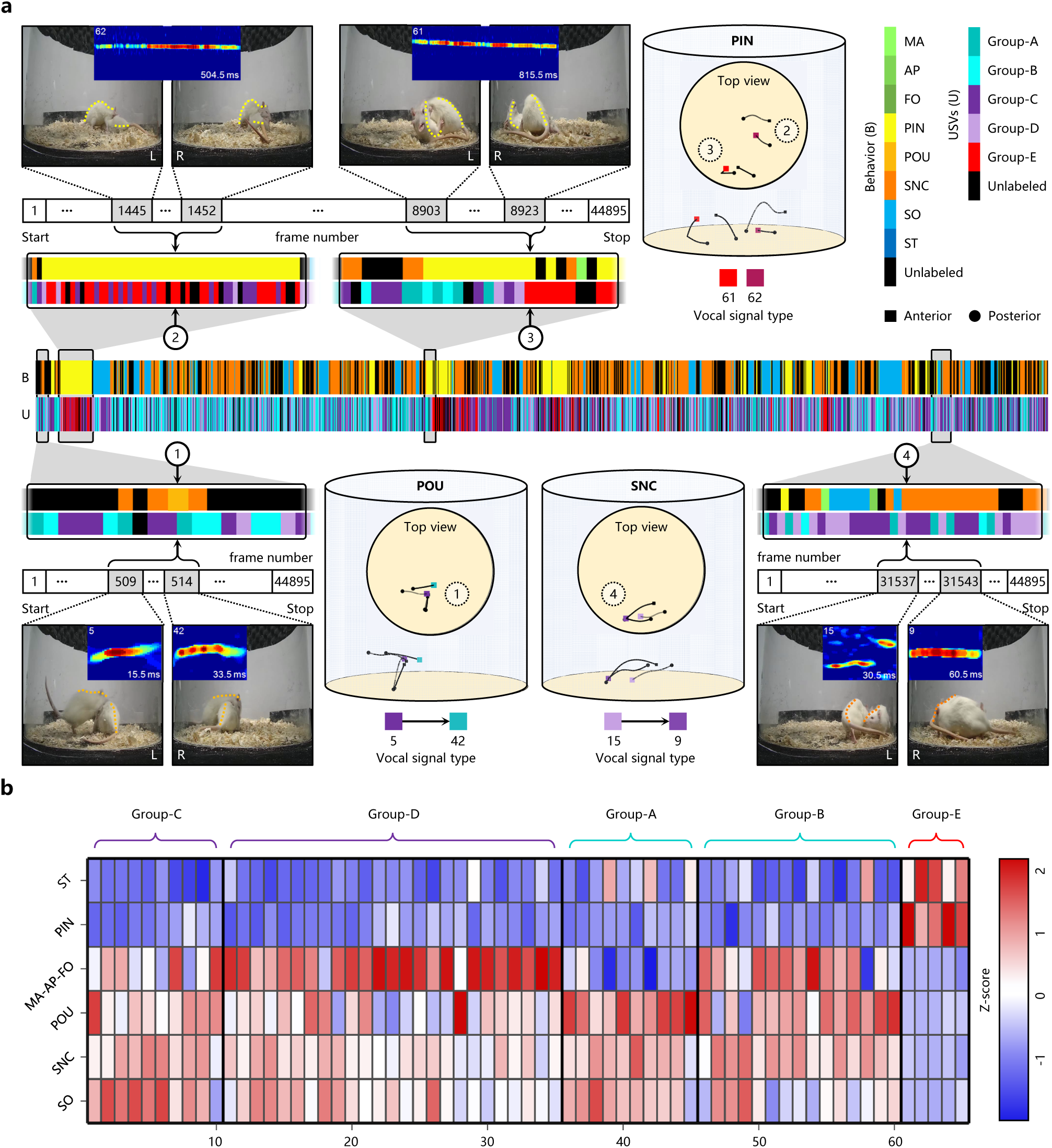
Revealing the behavior-vocalization interplay using ARBUR. **a**, Examples showing how 22-kHz USVs are associated with pinning (top), and appetitive 50-kHz USVs are associated with pouncing (bottom left) and social nose contact (bottom right). Frequency range of spectrograms: 20 kHz for clusters 5, 9, and 42; 0-40 kHz for clusters 61 and 62; and 40 kHz for cluster 15. **b**, Quantification of behavior-associated USV distributions, which indicates whether a certain cluster of USVs is emitted during a specific type of behavior above chance (red), at chance (white), or below chance (blue). Groups A-E, the same as in Figure 2. **Figure 4—figure supplement 1.** The behavior and USV ethograms of the simultaneously recorded vocal and behavioral repertoires.

The finding that during an aggressive interaction (i.e., pinning), the submissive one produces significantly more aversive 22-kHz USVs, is consistent with former research ***Burgdorf et al.*** (***2008***); ***Lukas and Wöhr*** (***2015***); ***Litvin et al.*** (***2007***). We also show that even though pouncing and pinning are similar types of behavior, they actually have distinct USV distributions. In particular, during pinning, the submissive rat rarely emits appetitive 50-kHz USVs, whereas, during pouncing, the submissive one produces mainly 50-kHz USVs. This may indicate that the submissive rat has distinct feelings and emotional states when engaging in the two types of behavior. ARBUR can reveal such a distinction because it can not only accurately discriminate these behaviors, but locate the submissive rat spatially. These findings are in line with former studies and therefore validate the effectiveness of ARBUR.

In addition, ARBUR reveals that several USVs, mostly 22-kHz USVs, are recorded when both rats are still (Figure 4b), indicating that some USVs may be emitted unconsciously (possibly during sleeping), and these 22-kHz USVs may neither convey emotional information nor trigger behavioral changes of the other rat. However, the still-associated (or sleeping-associated) USVs have rarely been focused on because of their independence from social interaction. Furthermore, ARBUR shows that rats produce more step 50-kHz USVs than non-step ones during motion (Figure 4b), indicating a part of step 50-kHz USVs may be caused by rats’ locomotion. Recent research has demonstrated the tight correlation between respiratory activity and frequency features of USVs ***Riede*** (***2013***), which indicated that locomotion may induce more step 50-kHz USVs by affecting rat respiratory activity.

## Discussion

Understanding the behavior-vocalization interplay of rats is inhibited by the difficulty of relating the behaviors and the USVs of freely behaving rats in complex social contexts. In this study, we propose a machine learning-based analysis system (named ARBUR) to relate rat vocalizations to their free behaviors. ARBUR contains three modules: 1) an unsupervised learning algorithm (AR-BUR:USV) that clusters rat USVs in an unbiased manner by comprehensively considering their mean frequency, duration, and both non-step (continuous) and step (non-continuous) contours in the spectrograms; 2) a machine learning-based hierarchical framework (ARBUR:Behavior) that detects the type of social behavior from a laterally binocular view; and 3) a sound source localization algorithm (ARBUR:SSL) that spatially allocates the USVs to the vocal rats.

Rats produce diverse USVs in terms of duration, mean frequency, and frequency contours ***Wright et al.*** (***2010***). Existing studies have only clustered the USVs in a way that considers some of these factors in an unsupervised ***Takahashi et al.*** (***2010***); ***Sangiamo et al.*** (***2020***) or manual ***Riede*** (***2013***); ***Brudzynski*** (***2013***) manner (Table 2). Among them, a recent study leveraged unsupervised learning to cluster mouse non-step USVs into 22 categories based on their contours, which showed potential for unbiased clustering of a broader spectrum of USVs ***Sangiamo et al.*** (***2020***). However, an unsupervised clustering algorithm that comprehensively takes all these three factors and step signals under consideration remains missing. To our knowledge, ARBUR:USV is the first unsupervised clustering method that considers the step USVs of rodents. Although it is designed and optimized to cluster rat USVs, it can be readily generalized for mouse USV applications. Compared with algorithms from existing studies, the proposed unsupervised clustering algorithm features: 1) the comprehensive consideration of frequency, duration, and contour information; and 2) the inclusion of both step and non-step USVs. Moreover, ARBUR:USV provides multiple choices for users to determine the optimal clusters, including arbitrary setting, the progressive method, and the elbow method. It should therefore be adaptive to a wide range of applications regarding rodent acoustic clustering and analysis.

**Table 2.** ARBUR:USV achieves comprehensive unsupervised clustering of rodent’s USVs compared with current research.

|  |  | **Manner of clustering** | **Mean frequency** | **Duration** | **Non-step contour** | **Step contour** | **Choice of optimal clusters (number)** | **Rodents** |
| --- | --- | --- | --- | --- | --- | --- | --- | --- |
| Wright et al. *Wright et al. (2010)* |  | Manual | ✓ | - | ✓ | ✓ | Arbitrary (15) | Rats |
| Riede et al. *Riede (2013)* |  | Manual | ✓ | - | ✓ | ✓ | Arbitrary (6) | Rats |
| Burgdorf et al. *Burgdorf et al. (2008)* |  | Manual | ✓ | - | ✓ | ✓ | Arbitrary (3) | Rats |
| Fonseca et al. *Fonseca et al. (2021)* |  | Supervised | ✓ | - | ✓ | ✓ | Arbitrary (11) | Mice |
| Sangiamo et al. *Sangiamo et al. (2020)* |  | Unsupervised | - | - | ✓ | - | Progressive (22) | Mice |
| Takahashi et al. *Takahashi et al. (2010)* |  | Unsupervised | ✓ | ✓ | - | - | Arbitrary (3) | Rats |
| MUPET *Van Segbroeck et al. (2017)* |  | Unsupervised | - | ✓ | ✓ | - | Arbitrary (20-200) | Mice |
| DeepSqueak *Coffey et al. (2019)* |  | Unsupervised | - | ✓ | ✓ | - | Automatic (20) | Mice |
| AVA *Goffinet et al. (2021)* |  | Unsupervised | - | ✓ | ✓ | - | Arbitrary (-) | Mice |
| **ARBUR:USV** |  | **Unsupervised** | **✓** | **✓** | **✓** | **✓** | **Multiple choices (65)** | **Rats** |

ARBUR:Behavior is a hierarchically supervised learning algorithm that archives state-of-the-art performance in classifying eight types of common rat behaviors, including three easy-to-confuse social behaviors (pinning, pouncing, and social nose contact). We show that the mainstream deep learning classification algorithms ***Touvron et al.*** (***2021***); ***Liu et al.*** (***2021***, ***2022***); ***He et al.*** (***2016***) are not competent for the end-to-end classification of rat complex behaviors. For example, ResNet achieves the best performance with an accuracy of only 0.333 using an eight-class classifier (Figure 3-figure supplement 2a). This highlights the necessity of hierarchical classification of rat complex behaviors. Therefore, ARBUR:Behavior hierarchically classifies rats’ non-social, moving, and social state behaviors sequentially. It detects non-social behaviors using the location feature of rats in the image and moving behaviors by estimating the optical flow of the rat as well as its relative orientation. To discriminate tricky social behaviors, ARBUR:Behavior extracts “histogram of oriented gradients” (HOG) ***Dalal and Triggs*** (***2005***) descriptors to represent the statistics of social posture features in a standardized way. This empowers ARBUR:Behavior with high-accuracy discrimination of rat social behaviors. ARBUR:Behavior outperforms existing studies with respect to comprehensive detection performance (Table 3).

**Table 3.** ARBUR:Bheavior achieves comprehensive behavior classification.

|  | Input modality | Algorithm characteristics | Background subtraction | Social behavior | Behavior categories | F1-score/<br>Accuracy |
| --- | --- | --- | --- | --- | --- | --- |
| DeepEthogram *Bohnslav et al. (2021)* | RGB (top view) | End-to-end (supervised) | - | ✓ | 6 | 0.638/- |
| SIPEC:BehaveNet *Marks et al. (2022)* | RGB (top view) | End-to-end (supervised) | ✓ | - | 3 | 0.720/- |
| DeepAction *Harris et al. (2023)* | RGB (top view) | Hand-crafted (supervised) | - | ✓ | 12 | -/0.739 |
| VSAMBR *Jhuang et al. (2010)* | RGB (side view) | Hand-crafted (supervised) | ✓ | - | 8 | -/0.771 |
| DeepCaT-z *Gerós et al. (2022)* | RGB-D (top view) | End-to-end (supervised) | ✓ | - | 4 | 0.822/- |
| OpenLabCluster *Li et al. (2022a)* | Pose and tracking data | Hand-crafted (semi-supervised) | - | - | 8 | -/0.842 |
| MARS *Segalin et al. (2021)* | RGB (top view) | Hand-crafted (supervised) | - | ✓ | 3 | 0.878/- |
| B-SOiD *Hsu and Yttri (2021)* | Pose and tracking data | Hand-crafted (unsupervised) | - | - | 11 | -/0.915 |
| **ARBUR:Behavior** | **RGB (side view)** | **Hand-crafted (supervised)** | **✓** | **✓** | **8** | **0.868/0.870** |

**Table 4.**
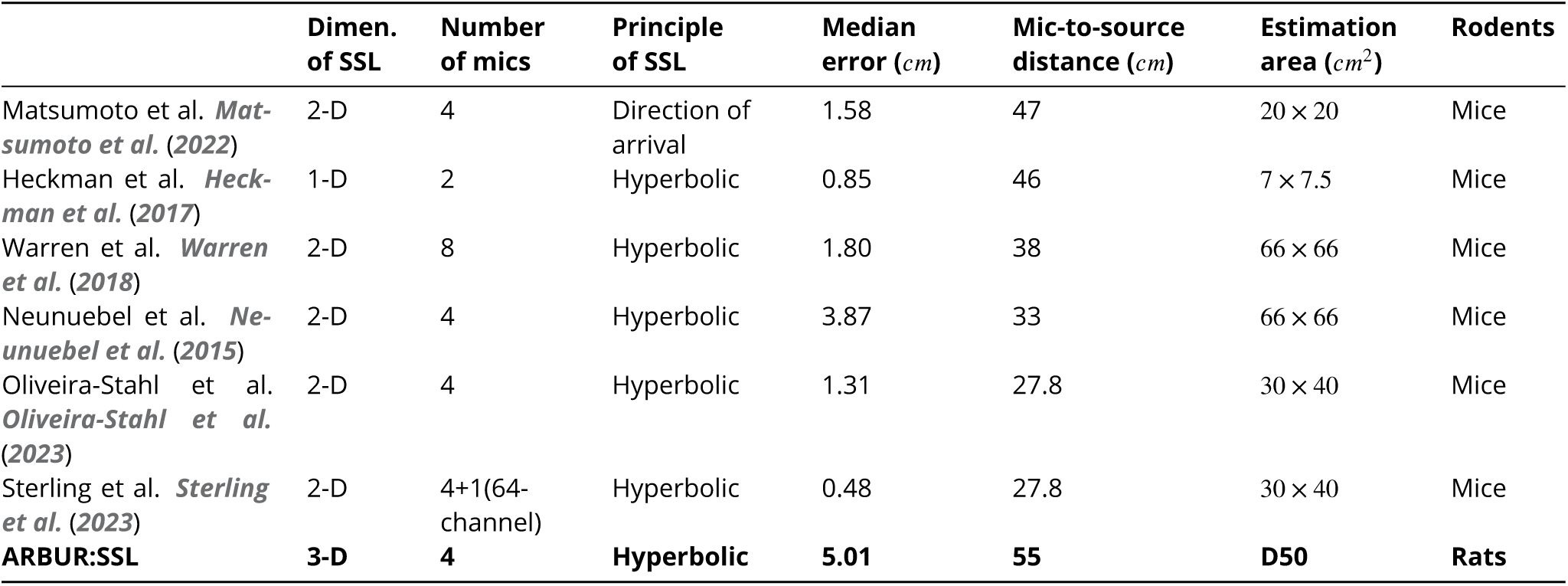
Comparison of ARBUR:SSL with mainstream sound source localization methods for rodents’ USVs.

ARBUR:SSL can locate the vocal rat during two-rat free-behaving scenarios by incorporating the lateral binocular view (for rat nose reconstruction in the Cartesian space), behavior classification results, height-sensitive distributed microphone configuration, and 3-D probabilistic triangulation of sound source. The large rat-to-mic distance in rat-rat interacting scenarios makes the recorded USV signal intensity (therefore signal-to-noise ratio) much weaker than others. ARBUR:SSL addressed this challenge by measuring the consistency between 3-mic SSL and 4-mic SSL results, hence ruling out those low-SNR USVs. It achieves desirable SSL precision suitable for rat-rat social scenarios. Another concern that haunted the experts working on SSL is the lack of accurate test datasets. Maintream studies reportedly induced mouse USVs using heterosexual urine ***Sterling et al.*** (***2023***); ***Oliveira-Stahl et al.*** (***2023***), but the visual localization of rodent snout would bring un-necessary errors, and lack the ability of snout height detection. In this work, we constructed a rat USV test dataset for evaluating SSL errors using USVs collected from still rats, which brings two advantages: 1) all the USVs are produced at the same spatial source position because the rat kept still throughout the process; 2) 3D position of the vocal rat nose/snout is provided, therefore allowing evaluating 3D SSL algorithms. ARBUR:SSL can therefore contribute to SSL research by highlighting the potential of refining low-SNR vocal signals and stressing the value of USVs emitted by still or sleeping rats for evaluating SSL algorithms.

By seamlessly combining these three modules, ARBUR features the advantages of comprehensive and unbiased USV clustering, hierarchical high-accuracy rat behavior detection, and 3-D sound source localization to reveal the latent behavior-vocalization interplay of rats. For example, during a socio-aggressive interaction (i.e., pinning), the submissive one produces significantly more aversive 22-kHz USVs, which is consistent with former research ***Burgdorf et al.*** (***2008***); ***Lukas and Wöhr*** (***2015***); ***Litvin et al.*** (***2007***). We also show that even though pouncing and pinning are similar types of behavior, they actually have distinct USV distributions. Moreover, ARBUR indicates several novel findings about still-associated or moving-associated USVs ***Riede*** (***2013***), which have long been neglected in the manual analysis-dominated research, and call for more stringent verification to bring substantial knowledge. We also note that although ARBUR is designed and optimized to investigate the behavior-vocalization interplay of rats, it can be potentially generalized to other rodents that also communicate through both non-verbal signals and USVs (for example, mice and mole rats).

Despite these advancements, ARBUR can be further improved in the following aspects. First, ARBUR:USV rejects the concurrent USVs (simultaneously recorded USVs at different frequencies), which may be important during social communication and could be separated by intelligent segmentation algorithms combined with high-precision SSL, further enriching the vocal repertoire. Moreover, ARBUR:Behavior lacks the ability to detect potential behavioral changes in freely behaving rats, which is essential for discovering the underlying mechanisms of novel behavior-vocalization and could be enhanced by incorporating unsupervised learning to uncover latent structures and categories in behavioral space ***Brattoli et al.*** (***2021***); ***Huang et al.*** (***2021***). In addition, ARBUR:SSL could be improved in terms of the limited number of visual reconstructions of rat noses by incorporating more camera views in the future, fulfilling the need for a more reliable three-dimensional sound source localization. Using ARBUR, we revealed that rat USV distribution is biased by rat behavior, indicating that the USVs of rats are associated with their behaviors. For example, a submissive rat significantly up-regulates the aversive 22-kHz USVs during pinning. Moreover, during pouncing and social nose contact, both rats produce substantially more appetitive 50-kHz USVs than they do during pinning.

In summary, we proposed a machine learning-based analysis system, which can not only automatically reveal the well-understood behavior-associated vocalizations that were carefully concluded by other behavioral researchers, but also hold the promise to indicate novel findings that can be hardly found by manual analysis, especially regarding step USVs and the active/passive rat-associated USVs during easy-to-confuse social behaviors. This work highlights the potential of machine learning algorithms in automatic animal behavioral and acoustic analysis and could help mechanistically understand the interactive influence between the behaviors and USVs of rats.

## Methods

### Animals and Experiment preparation

#### Animals

Adult (8-10 weeks) male (n = 4) and female (n = 4) Sprague-Dawley rats (stock number 198499, 212822, SPF biotechnology, Beijing, China) were used. Rats were kept on a 12/12 h light/dark cycle at a consistent ambient temperature (26±1^◦^C) and humidity (50±5%), and all experiments were performed during the light cycle. Food and water were accessed ad libitum. All experimental procedures were approved by the Institutional Animal Care and Use Committee of the Beijing Institute of Technology, Beijing, China. Before all recording experiments, rats were singly housed for at least 3 days to minimize group housing effects on the social behavior of the rats.

#### Video and audio stream recording

The video and audio simultaneous recording experimental set-up is shown in Figure 1–figure supplement 1. Rats were able to behave freely in a circular open-field arena made of a transparent acrylic barrel with a base diameter of 0.5 m and a height of 0.5 m. A black velvet cloth was placed underneath the recording area as well as wood chips to avoid reflecting light. Encircling the recording area with sound-absorbing material above can reduce sound reflections from the walls. Two synchronized cameras were mounted at an angle of 60 degrees on either side of the acquisition area. Four ultrasonic microphones (CM16/CMPA, Avisoft Bioacoustics, Glienicke, Germany) are fixed evenly to the edges of the transparent acrylic barrel. The video stream was recorded at 25 frames per second. The four channels of the audio stream from the four microphones were simultaneously recorded through data acquisition equipment (UltraSoundGate 416H, Avisoft Bioacoustics). The video and audio streams were synchronized by aligning the recording onset timing. Both the streams were stored on a high-performance computer to avoid recording time shifting (32G RAM). The video and audio stream recording experiments were conducted over 4 days (12 hours of continuous recording per day) with no external human interference.

### Audio segmentation

Audio streams were segmented automatically with multi-taper spectral analysis. The vocal signals from microphone 1 were segmented using the USVSEG algorithm ***Tachibana et al.*** (***2020***); ***Matsumoto et al.*** (***2022***), and the parameters of the algorithm were optimized as follows. First, multitaper spectrograms (time-frequency matrices) were generated using six discrete prolate spheroidal sequences of length 512 as windowing functions (NW=3) ***Percival and Walden*** (***1993***). Then the spectrograms were flattened by replacing the first three cepstral coefficients with zero to reduce transient broadband noises and subtracting the median spectrum. After flattening, we thresholded the flattened spectrograms with a threshold of 4.5 to further reduce the environmental noise. These spectrograms were band-pass filtered between 10 and 100 kHz. In addition, the maximum duration of an audio segment was set to 5000 ms and the minimum duration was set to 5 ms. Sound elements with a more than 30 ms gap were judged as two individual syllables and segmented as two independent spectrograms accordingly. Finally, these segmentation timings from microphone 1 were used to segment vocal signals from the other three microphones. A total of 43357 ultrasonic signals were segmented from all the audio data collected.

### Vocal signals clustering

#### Extraction of frequency contours

The segmented spectrograms (time-frequency matrices) were filtered sequentially by the median, Gaussian, and threshold filters to enhance the signal-to-noise ratio. The filtered spectrograms were then binarized for detecting the boundaries of the frequency contours (edges in the spectrograms) by the modified Moore’s neighbor tracking algorithm with the Jacobian’s termination condition. Those contours with less than 22 points and a maximum intensity inside the contour of less than 15 were detected as isolated islands and eliminated. For the discontinuous USVs with multiple contours, only the longest four contours were selected as the final contours to recapitulate the original complex signals. Harmonics were considered as those frequency contours with 90% overlap in time, and only the frequency contour of the lowest mean frequency was preserved whereas others were eliminated. Other than harmonics, those frequency contours that share an overlap of duration over 3 ms were considered overlapping vocalizations emitted by two rats and also abolished.

#### Coarse clustering

Coarse clustering consists of two steps. In the first step, the USVs were clustered coarsely into 2 groups according to their mean frequency: aversive 22-kHz and appetitive 50-kHz USVs. In the second step, the 2 groups were further divided into 5 groups (A-D) according to their distribution of duration and mean frequency of frequency contours: so-called 22-kHz, low-peak-frequency/high-peak-frequency step/non-step 50-kHz USVs (Figure 2–figure supplement 1a). Specifically, those USVs with a mean frequency of less than 32 kHz were classified as 22-kHz signals. The rest of the USVs (50-kHz) were clustered into 2 groups (low-peak-frequency 50-kHz and high-peak-frequency 50-kHz) by k-means clustering according to two features: duration and mean frequency. These USVs were further classified into 4 groups according to their continuity of frequency contours: non-step (continuous) USVs containing only one contour and step USVs containing more than one contour.

#### Determining the optimal number of clusters

In the third step, groups A-E were further divided using unsupervised learning within their groups. For the appropriate setting of the cluster number, ARBUR:USV provides users with three choices: arbitrary, automatic, and progressive methods. The arbitrary method allows users to set an integer number for each cluster. For example, in Figure 2, cluster numbers for groups A-E are set as 10, 25, 10, 15, and 5, respectively. The automatic method relies on the well-known elbow method to avoid over-clustering, which works as follows. For each cluster number, the total within-cluster error (TWCE) is calculated. By sliding from 2 clusters to 100 clusters (200 for group B for its complexity), forming a TWCE-clusters curve. Then, for each point on this curve, its left and right sides are used for linear fitting, and the total fitting error is stored, which forms a second curve: fitting error versus cluster number (see Figure 2–figure supplement 2 as an example). The global bottom of the second curve is called the inflection point, which indicates the optimal number of clusters. The progressive method takes a user-defined intra-cluster variance parameter to find the appropriate number. It calculates the innerpoint percentage (percent of points falling within the mean±2.5SD range of each cluster). Theoretically, with increasing clusters, the intra-cluster variance decreases. This method starts with 2 clusters, calculates the average innerpoint percent, and stops when the user-defined threshold is first reached. For example, if the threshold is set as 98%, that is, an average of 98% points should be the inner points for all clusters and all features. The optimal cluster should be 8 for group E (see Figure 2–figure supplement 3).

#### Refined clustering of non-step USVs

The non-step (continuous) signals were clustered further according to their contour shapes. The detected boundary points were centered by subtracting the mean frequency. To normalize these centered contours with different durations, we constructed a fixed-length feature vector containing 100 features (Figure 2–figure supplement 1b). The first 50 and last 50 features represent the upper and the bottom boundaries of the centered contour, respectively. Each contour was linearly mapped to a 100-feature vector. By doing so, frequency contours with different durations or mean frequencies can be clustered appropriately according to their shapes. These USVs were categorized using k-means clustering based on the 100-feature vectors. The number of clusters for low-frequency and high-frequency USVs were both arbitrarily set as 10. Alternatively, the number of clusters can be determined progressively as in ***Sangiamo et al.*** (***2020***).

#### Refined clustering of step signals

For the step (discontinuous) signals with multiple contours, a similar 100-feature vector was constructed based on these contours (Figure 2–figure supplement 1b). All contours of each USV were centered by subtracting their mean frequencies. Then, the upper boundaries of each contour were sorted in ascending order in time and mapped linearly to a 50-feature vector. By doing so, those inter-contour intervals were deleted, and contours were placed compactly. Similarly, the bottom boundaries were mapped to the second 50-feature vector. These step USVs were then categorized using k-means clustering based on the 100-feature vectors concatenating these two vectors together. The numbers of clusters for low-frequency and highfrequency USVs were arbitrarily set as 15 and 25, respectively.

#### Refined clustering of 22-kHz USVs

22-kHz USVs vary a lot in duration, ranging from several microseconds to several seconds, such a feature should not be ignored in clustering. In contrast, the contour shapes varied little across USVs. Therefore, we constructed a 2-feature vector for each 22-kHz signal containing the mean frequency and duration. These 22-kHz signals were then categorized using k-means clustering based on the 2-feature vectors. The number of clusters was arbitrarily set to 5.

#### Validating clustering

To validate clustering results, dissimilarities within each cluster (intra-cluster variance) and between different clusters (inter-cluster distance) were evaluated, respectively. They were calculated based on the constructed 100-feature vectors for both step and non-step USVs (Figure 2–figure supplement 5a). The inter-cluster distance of the *i*th cluster *D*_*i*_ was defined as the average difference between each pair of frequency contours from two different clusters within each group (one of 5 groups by coarse clustering):

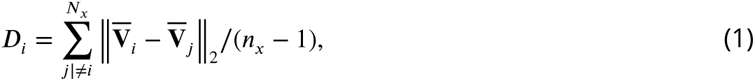

where ‖⋅‖_2_ denotes *l*_2_-norm, *N*_*x*_ is the index integer set of one of five particular groups, 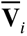 ∈ *R*^100^ is the average feature vector of the *i*th cluster in that group and *n*_*x*_ is the number of individuals in *N*_*x*_. The intra-cluster variance of *i*th cluster VAR_*i*_ was calculated as:

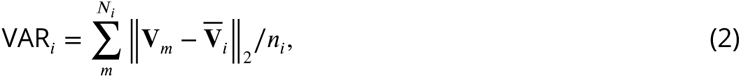

where *N*_*i*_ is the index integer set of the *i*th cluster, **V**_*m*_ is the feature vector of the *m*th USV in this cluster, and *n*_*i*_ is the number of individuals in *N*_*i*_.

### Behavior classification

The workflow of ARBUR: Behavior is shown in Figure 3–figure supplement 1. ARBUR:Behavior uses the binocular side-view video stream (segmented by SegFormer to remove the background) as input to discriminate eight behaviors of rats. First, the rat regions within each input image were labeled by edge detection. Detected regions were recognized as outlier regions if their areas were below the maximum error range:

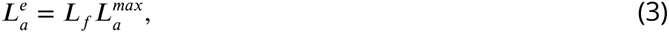

where 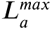 is the size of the maximum area with labels in the current image, and *L*_*f*_ is the filter threshold. The segmented image and labels were then input into a hierarchical algorithm to classify the non-social, moving, and social state behaviors of rats as follows.

#### Non-social state behaviors

The movement variation (in pixels) *M*_*p*_ of the center position of each label was calculated to determine whether rats have moved:

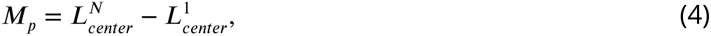

Where 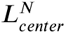 and 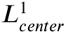 are the center pixel positions of the labels in the last and the first image. The value of *M*_*p*_ for sleeping rats hardly changes. For individually moving rats, we consider the total number of labels in each image to reflect whether the rats come into contact with each other (solitary) within that sequence of images.

#### Moving state behaviors

Based on the behavioral definition in Table 1, we used the number of labels in the first and last images of the image sequence to initially determine the moving state behaviors. Then, we estimated the motion optical flow of the rat in the images and defined its centroid motion vector as:

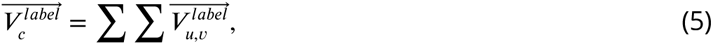

where *u* and *v* are the pixel coordinates of the image. As shown in the moving state behaviors section of Figure 2–figure supplement 1, we determined the specific behavior by judging the vector 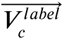 relative direction of the two labels.

#### Social state behaviors

We designed a decision tree algorithm based on multiple binary SVM classifiers to detect social state behaviors. A total of 860 images were labeled (PIN: 280, POU: 300, SNC: 280), and the HOG features were extracted as the training dataset. The SVM models were trained using a third-degree polynomial kernel function, and parameter optimization was performed for each model. Specifically, for each image, we deployed three SVM models simultaneously (PIN-POU, PIN-SNC, POU-SNC) to determine the category of the image with maximum probability. We defined the total confidence of the behavioral category for all images:

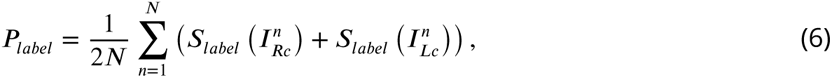

where

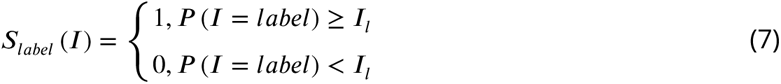

where *P* (*I* = *label*) is the probability that the current image category is *label* (PIN, POU, SNC), and *I*_*l*_ is the likelihood of a specific image belonging to the category. Behavioral category *I*_*c*_ was mapped through the maximum probability in each *P*_*label*_ :

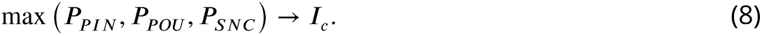

Image sequences that do not meet the aforementioned criteria were labeled as undetected.

### Sound source localization

#### 3D keypoints detection of rats

To obtain the spatial position of the two rats, we designed a deep learning-based keypoint detection network capable of detecting seven keypoints distributed across the rats’ heads, spines, and tails. We then calculated the 3D coordinates of the keypoints via triangulation by utilizing the calibrated camera parameters and the detected key points in two image spaces. It is worth noting that for rats engaged in social behaviors, there can be occlusions of certain body parts, resulting in only a part of keypoints being detectable in each frame. Using these 3D coordinates, a curve that represents the posture of the rat in the current frame could be fitted (Figure 4a), which could be useful to determine the submissive rat and the dominant one during PIN (pinning) and POU (pouncing). For other cases, only the 3D coordinates of the rats’ noses were used for the downstream SSL processes.

#### Estimating sound sources

The sources of USVs were estimated using an algorithm modified from Neunuebel et al. ***Neunuebel et al.*** (***2015***). The workflow of this algorithm is illustrated in Extended Data Fig.5. For each extracted USV from four microphones, a data augmentation process was performed by slicing the recognized frequency contours into *m* segments. Then, four groups with 3 channels (mics) and a group with 4 mics were input to a steered response power-based sound source localization method. That is, *m* × 5 estimates were calculated. The estimates with 4 mics (*m* × 4) estimates and those with 3 mics (*m* × 5) were used to calculate two 2-D probability density functions (*f*_3_(*x*, *y*), *f*_4_(*x*, *y*), *x*, *y* ∈ ℝ), respectively. Then, the localization confidence index (LCI) *P*_*LCI*_ that represents the reliability of estimation results was produced as follows.

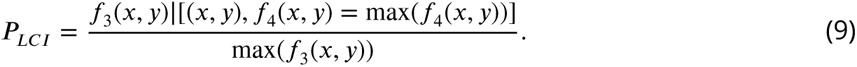

If the LCI is above the pre-set threshold (0.6), the SSL result is accepted as the center of estimates with maximum probability densities in *f*_3_(*x*, *y*) and *f*_4_(*x*, *y*). The SSL results are rejected otherwise.

#### Combining visual and SSL estimations

Because rats have a body length as long as 25 cm, the height of their sound source cannot be ignored in some cases. For example, if a rat presents a high-rearing pose, the nose can be as high as over 20 cm. Under these circumstances, the height should be considered to avoid incorrect SSL prediction.

For those behaviors with no apparent height difference between the noses of two rats (besides POU and PIN), the SSL results were assigned to rat noses based on horizontal information only. That is, the USV was assigned to the nearer rat if two noses were detected. If only one rat nose was detected, the USV was assigned to the detected one if it lies within the SSL estimation area (10 cm from the estimated point), and the USV was assigned to the undetected one otherwise. If no rat noses were detected, the USV cannot be assigned.

For pouncing and pinning behaviors, the SSL results were assigned based on not only horizontal information but also 3-D information. If two rat noses were detected, SSL results were calculated again with the nose height added. The USV was allocated to the rat with a higher LCI. If only one rat nose was detected and an obvious height difference was observed and labeled, the USV can be allocated to the rat with a higher lCI by assuming the height of another rat. This 3D SSL process is useful if two rats are close but have obvious height differences, which is common in pinning and pouncing.

### Statistics

All statistical analyses were performed in MATLAB (version 2021b, MathWorks) using two-sample single-tailed Welch’s t-tests with a significance level of *α* = 0.05. Data distributions were assumed to be normal, but this was not formally tested.

## Declarations

### Data Availability

All data needed to evaluate the conclusions in the paper are present in the Article. The datasets generated and/or analyzed during the current study are available from the corresponding author upon reasonable request.

### Code Availability

The open-source software ARBUR and its tutorials are freely available at GitHub (https://github.com/ Guanglu-Jia/ARBUR) and Zenodo (https://doi.org/10.5281/zenodo.8081539) ***Jia et al.*** (***2023***).

## Acknowledgments

This study was funded by the National Natural Science Foundation of China under Grant 62088101, the Science and Technology Innovation Program of Beijing Institute of Technology under Grant 2022CX01010, and the National Science and Technology Key major projects (STI2030-Major Projects 2022ZD02068000).

## Author contributions

Q.S. conceived and supervised the project. Z.C. wrote the manuscript and analyzed the results. Z.C. and G.J. implemented the methods, conducted the experiments, and processed the data. Q.Z. and Y.Z. assisted in conducting the experiments. All authors contributed to discussions. All authors edited and approved the manuscript.

---

**Figure 1—figure supplement 1.**
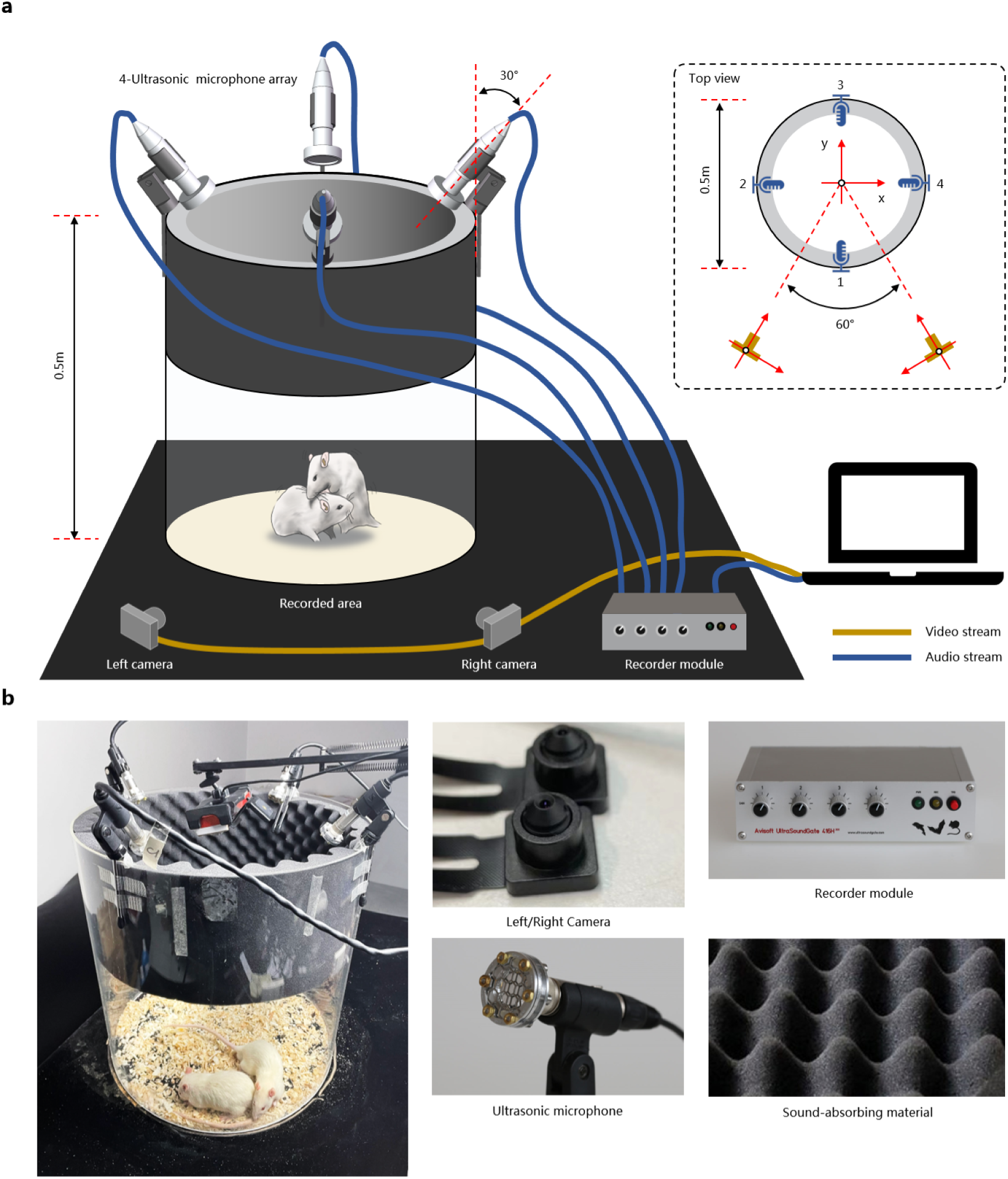
Experimental setup for recording video and audio streams of rats simultaneously. **a**, Schematic illustration of the setup. The video stream is recorded by two lateral cameras fixed on the ground, and the audio stream is recorded by a 4-mic array, which is fixed above the transparent acrylic arena. **b**, Images of the setup and devices.

**Figure 2—figure supplement 1.**
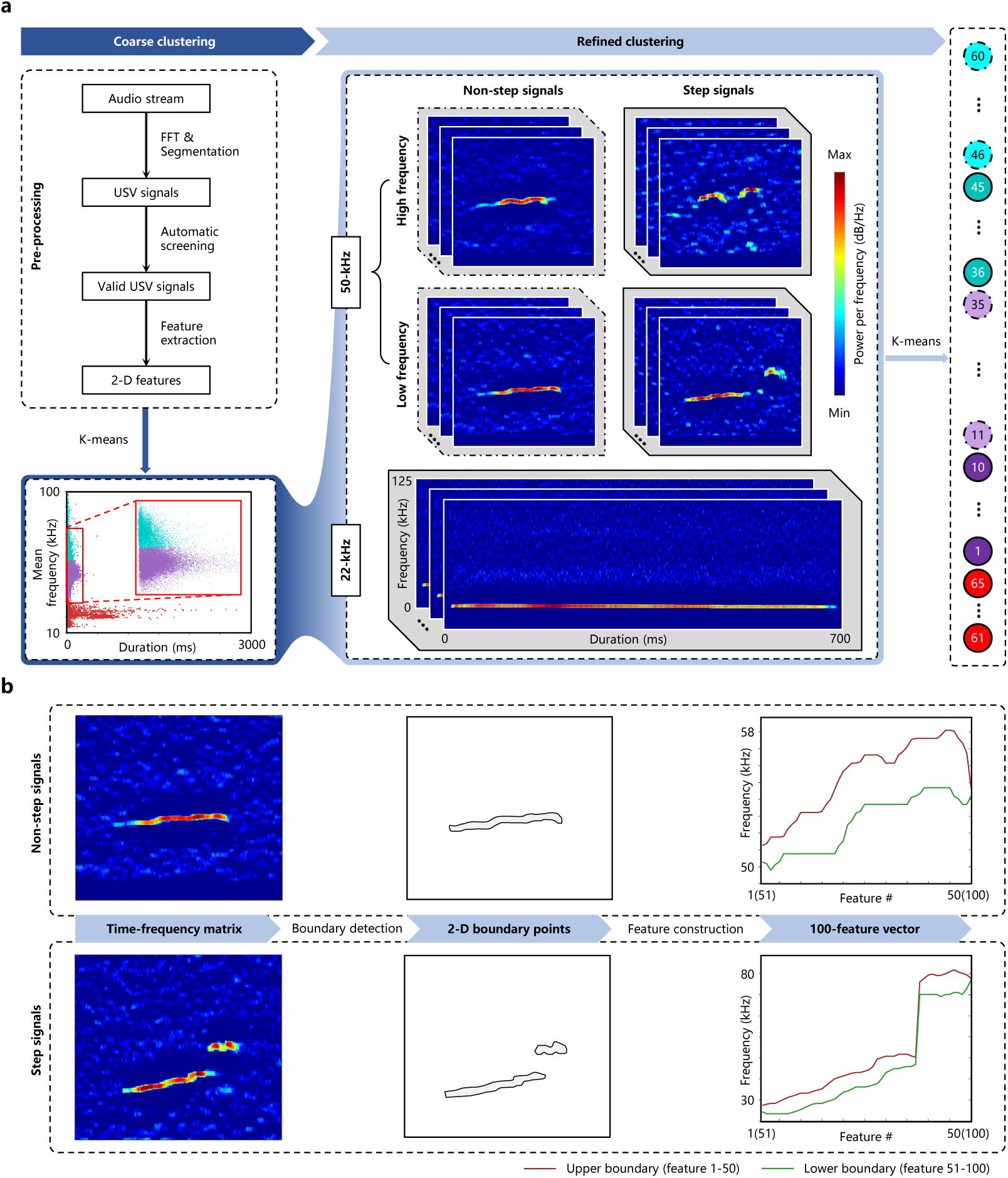
Workflow of the unsupervised clustering algorithm ARBUR:USV. **a**, In the first step (coarse clustering), the recorded audio stream was Fast Fourier Transformed, segmented, screened, and clustered into five groups sequentially; in the second step (refined clustering), USVs were clustered according to their frequency contours (for 50-kHz USVs) or mean frequency and duration (for 22-kHz USVs) within each group. **b**, Feature construction processes for non-step and step 50-kHz USVs.

**Figure 2—figure supplement 2.**
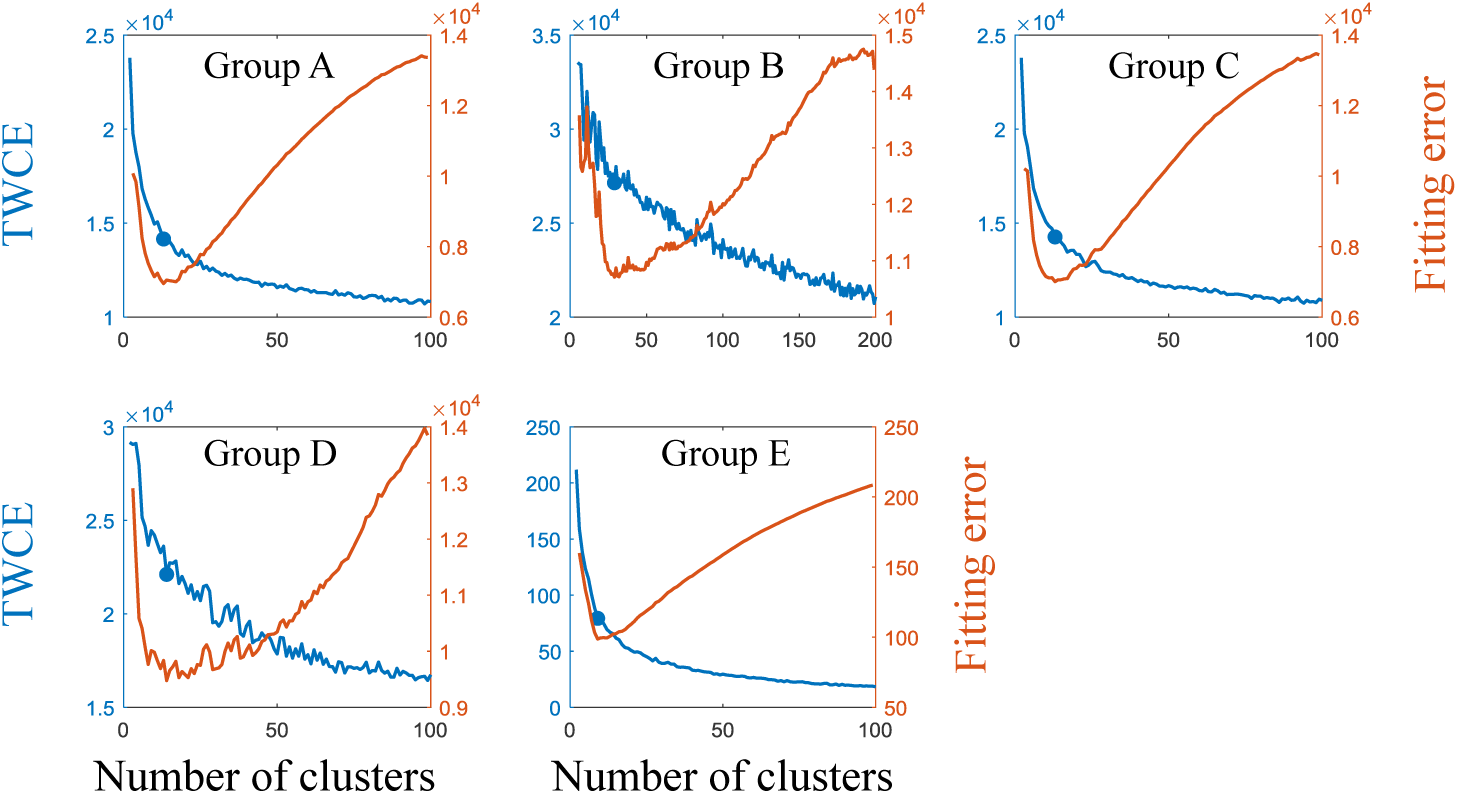
Determining the optimal number of clusters for groups A-E using the elbow method in our dataset. The optimal number for groups A-E are 13, 28, 13, 14, and 8, respectively. Left axis: total within-cluster error (TWCE); right axis: fitting error of two lines to the left and right side of a certain number. The lowest fitting error reveals the inflection point of the TWCE-number curve, which is the optimal number of clusters.

**Figure 2—figure supplement 3.**
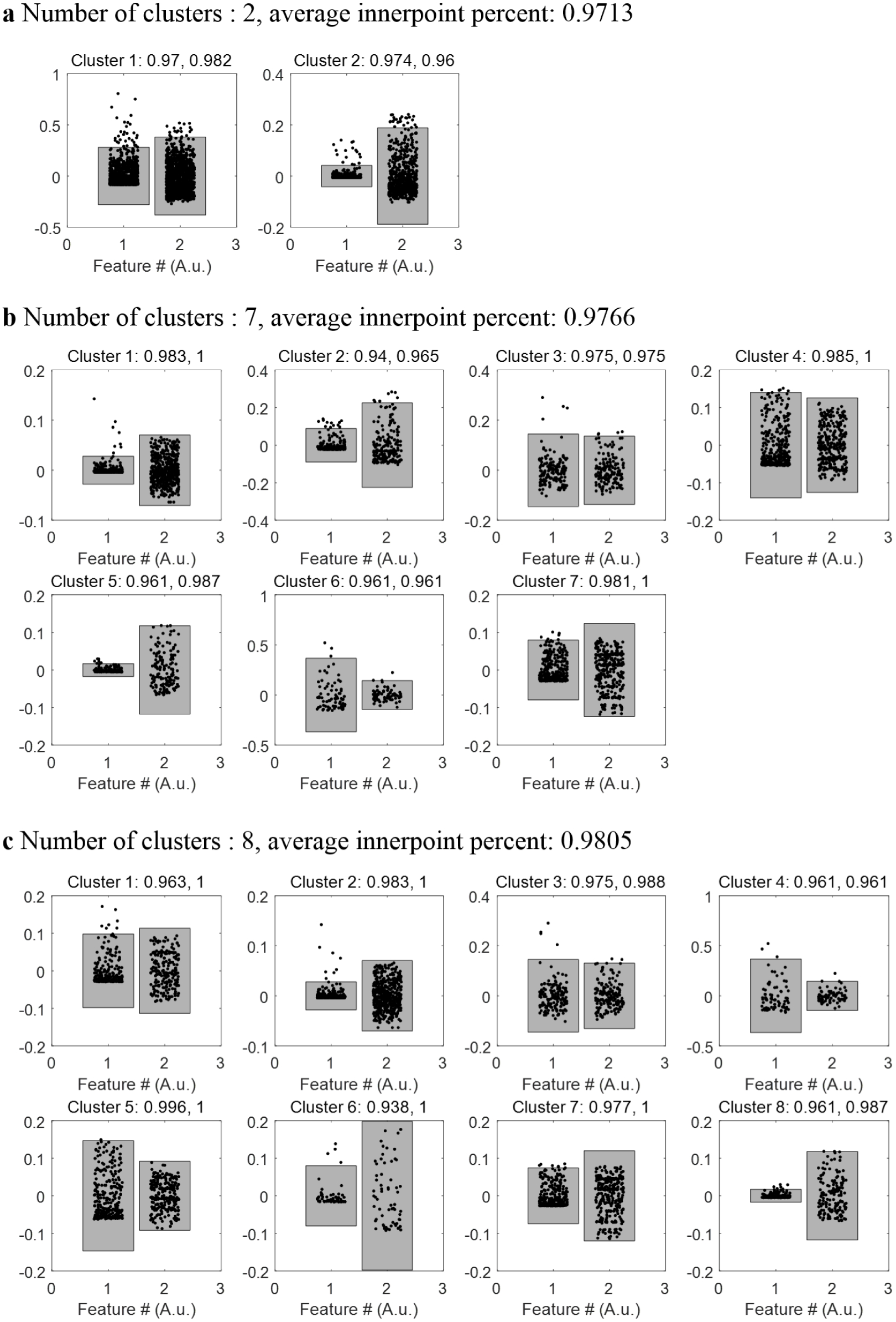
Determining the optimal number of clusters using the progressive method for group E. With increasing numbers, the intra-cluster variance gradually decreases, and the average innerpoint percent (within Mean ± 2.5 SD) increases. The progressive method starts with two clusters. When the threshold is set as 98%, it was first reached for cluster number = 8, hence the optimal number for 98% threshold is eight.

**Figure 2—figure supplement 4.**
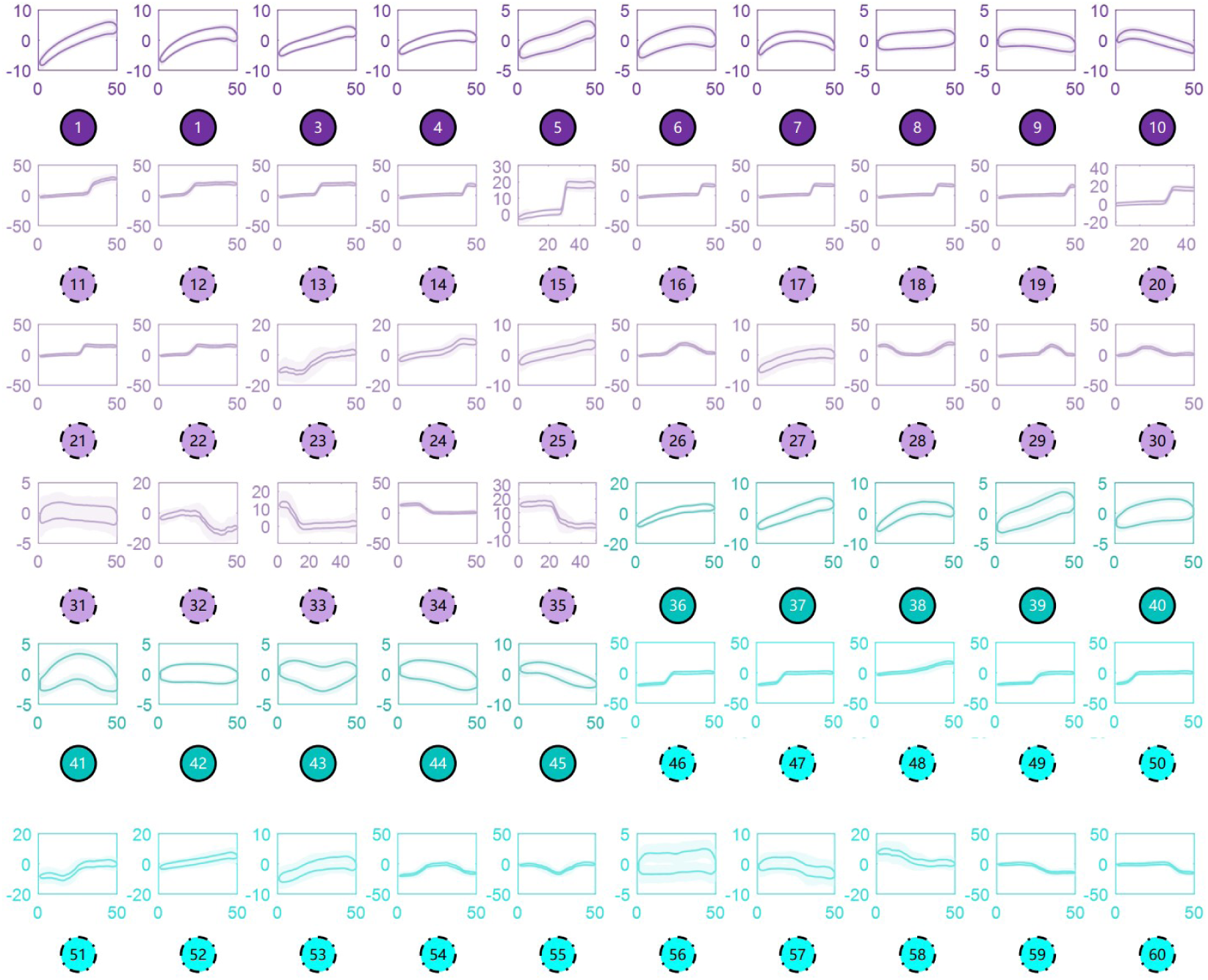
Mean value (solid lines) ± s.e.m. (shadow) of frequency contours within each cluster for clusters 1-60.

**Figure 2—figure supplement 5.**
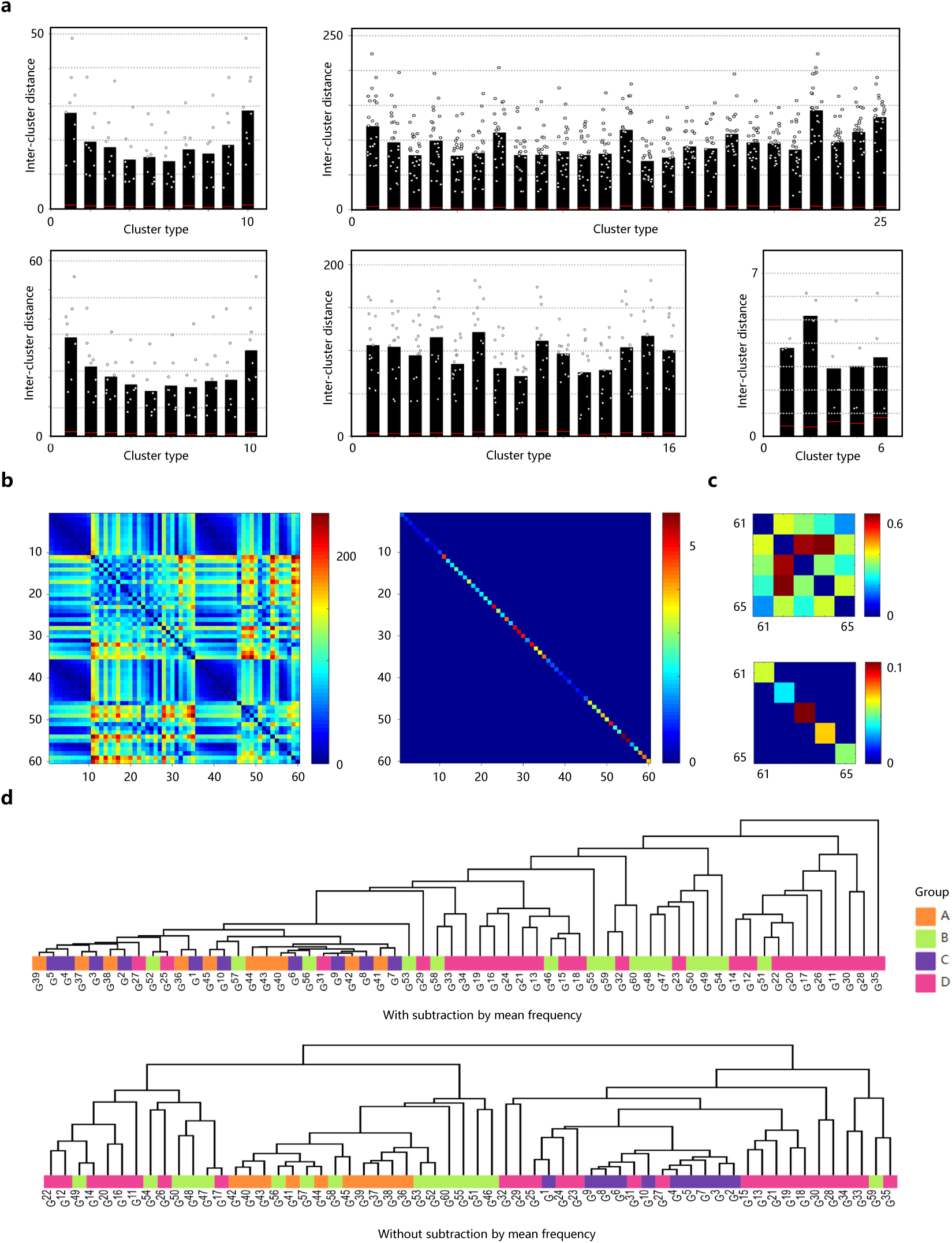
Validating ARBUR:USV. a, Inter-cluster distances between two clusters within each group, indicated by black circles. Black bars represent the average inter-cluster distance of this cluster with respect to the rest clusters within the same group. Intra-cluster variances of each cluster are indicated by red lines. b, Matrices of inter-cluster distance (left) and intra-cluster variance (right) for clusters 1-60. c, Matrices of inter-cluster distance (left) and intracluster variance (right) for clusters 61-65. d, Dendrograms of hierarchically clustered results for clusters 1-60 with (top) and without (bottom) feature vectors being subtracted by mean frequency.

**Figure 2—figure supplement 6.**
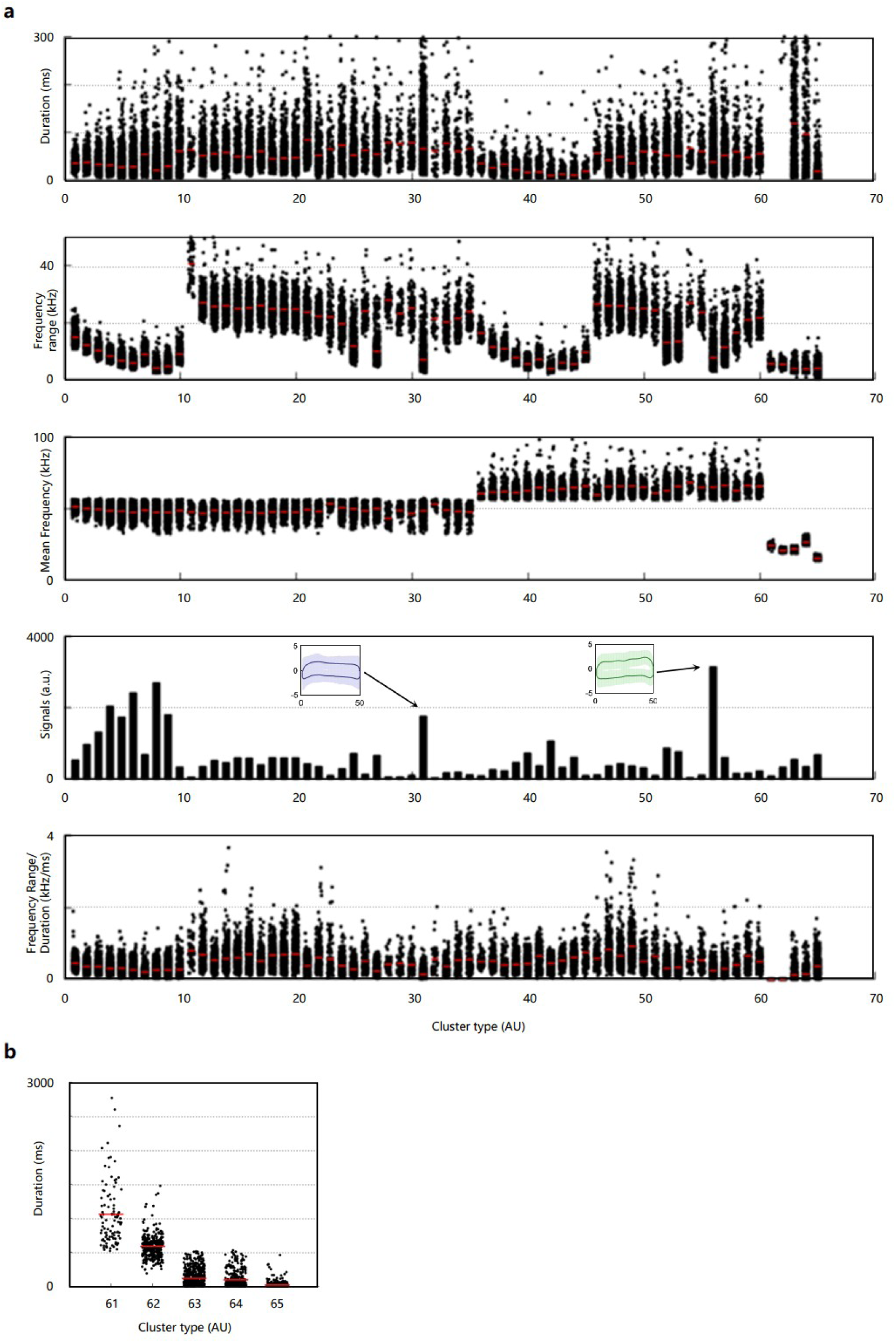
Quantification of the clustering results. a, The duration, frequency range, mean frequency, USVs in each cluster, and frequency range duration across clusters. b, Zoom-in view of the duration for clusters 61-65.

**Figure 3—figure supplement 1.**
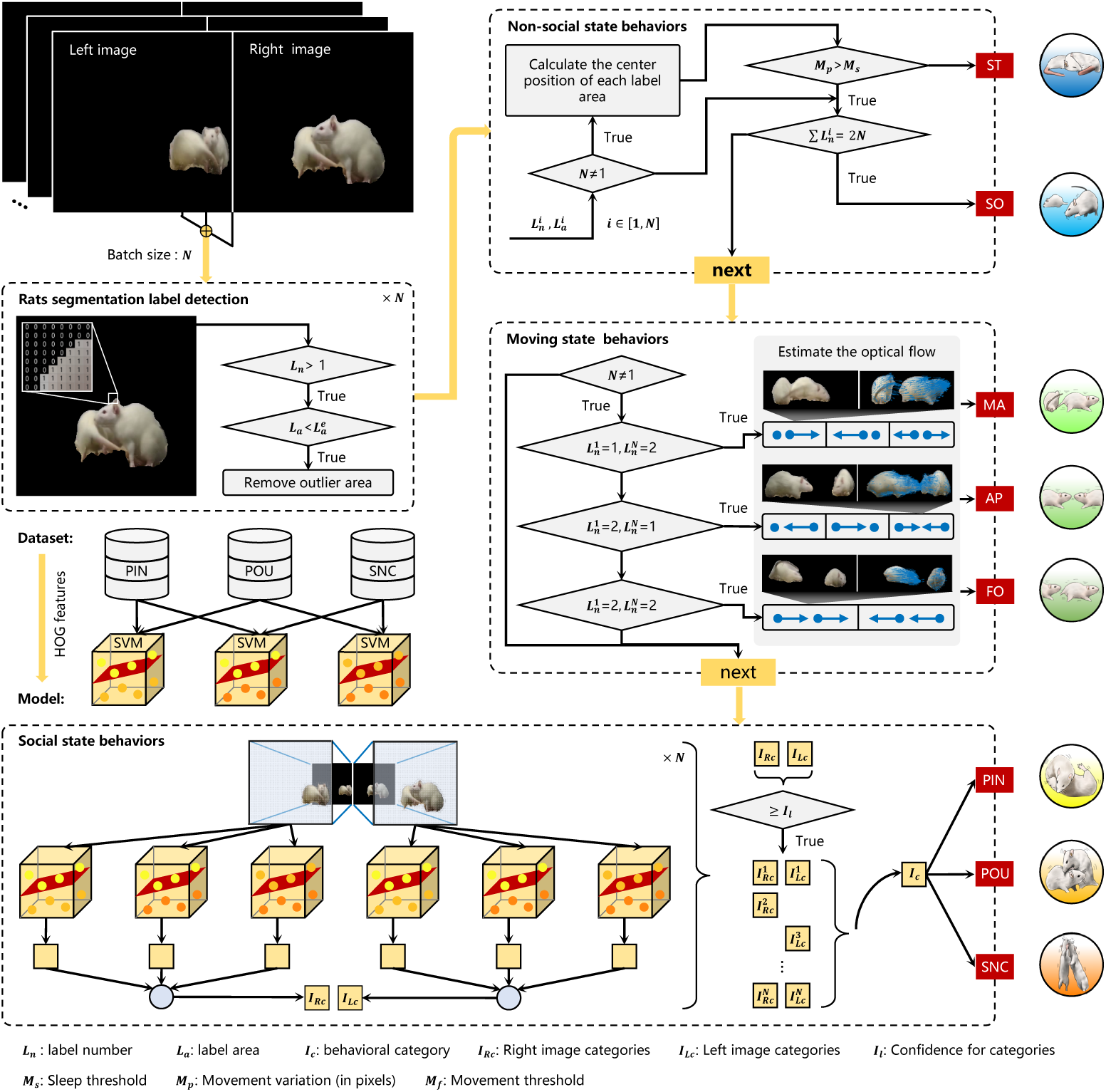
Workflow of ARBUR:Behavior. Rat regions (foreground) of the binocular image sequences (recognized by SegFormer) are labeled by edge detection. Both rat regions and labels are input to a hierarchical algorithm to classify rats’ non-social, moving, and social state behaviors sequentially. Non-social state and moving state behavior are detected by displacement and motion optical flow of the rats. A decision tree algorithm based on multiple binary SVM classifiers discriminates easy-to-confuse social state behaviors. ST: still; SO: solitary; MA: moving away; AP: approaching; FO: following; PIN: pinning; POU: pouncing; SNC: social nose contact.

**Figure 3—figure supplement 2.**
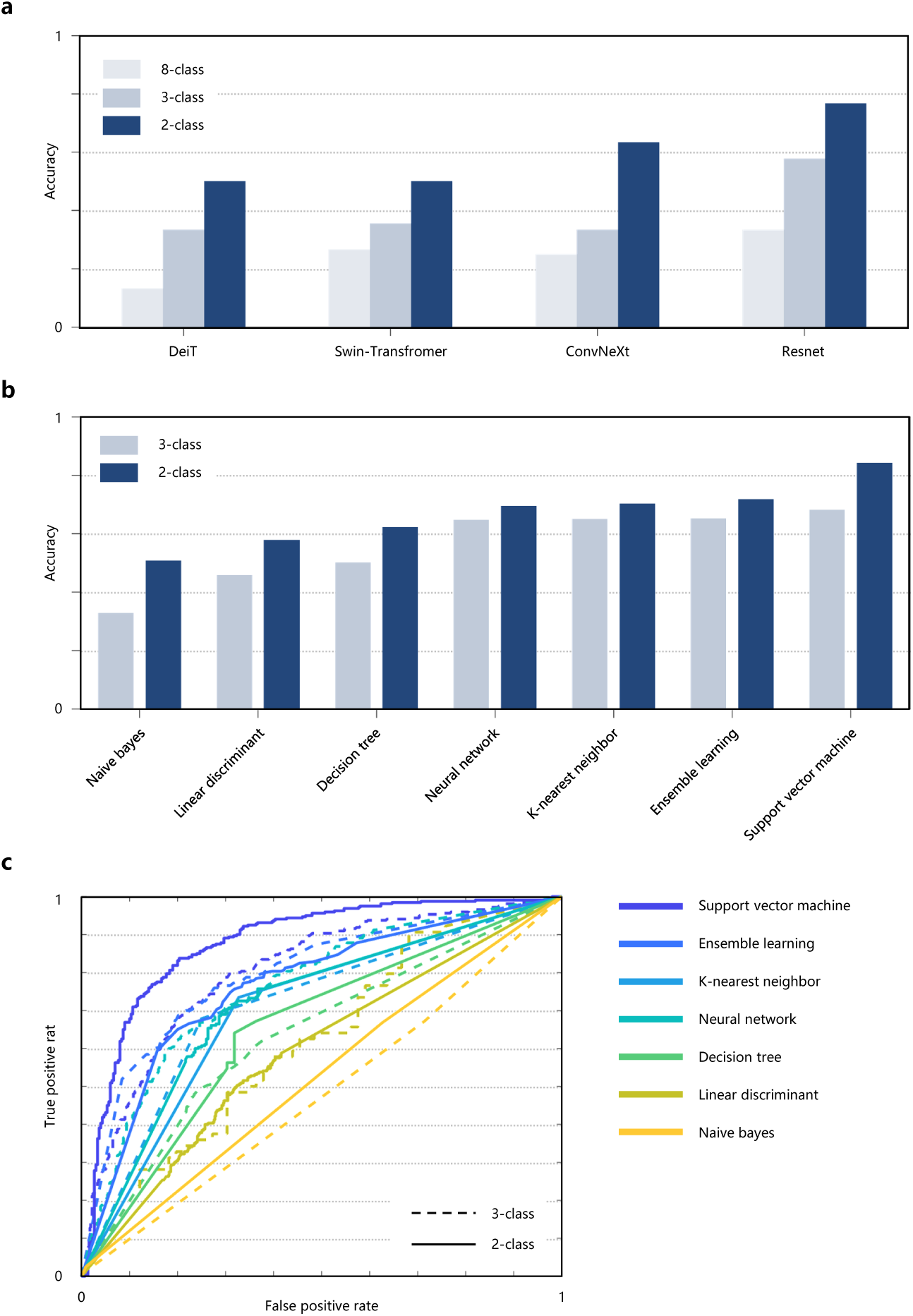
Comparison of algorithms for discriminating the easy-to-confuse social state behaviors. **a**, the average accuracies of four mainstream end-to-end classification networks for detecting 2-class, 3-class, and 8-class behaviors (including all behaviors), respectively. **b**, the average accuracies of seven classic machine learning-based algorithms for 2-class and 3-class behavior discrimination. **c**, The ROC (receiver operating characteristic) curves of the seven algorithms for 2-class and 3-class behavior discrimination.

**Figure 3—figure supplement 3.**
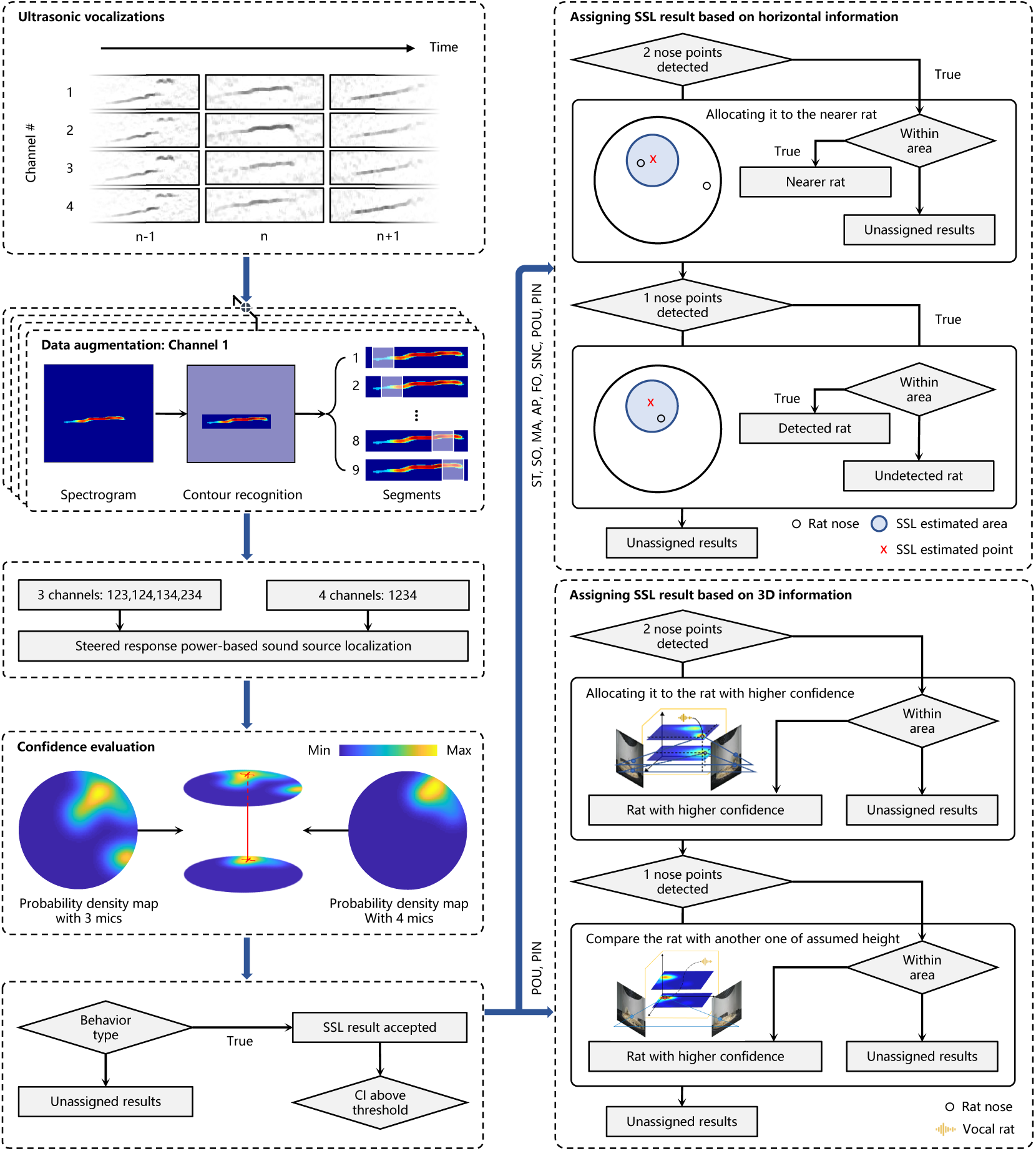
Workflow of ARBUR:SSL. Segmented ultrasonic vocalizations (USVs) from four channels (mics) were augmented, assembled (with 3 or 4 channels), and input into a steered response power-based sound source localization algorithm. Then, the localization confidence indices (LCIs) were evaluated and the SSL results were calculated, which were combined with 3D coordinates of rats’ noses to assign the USVs to the vocal rats by either horizontal or 3D information.

**Figure 3—figure supplement 4.**
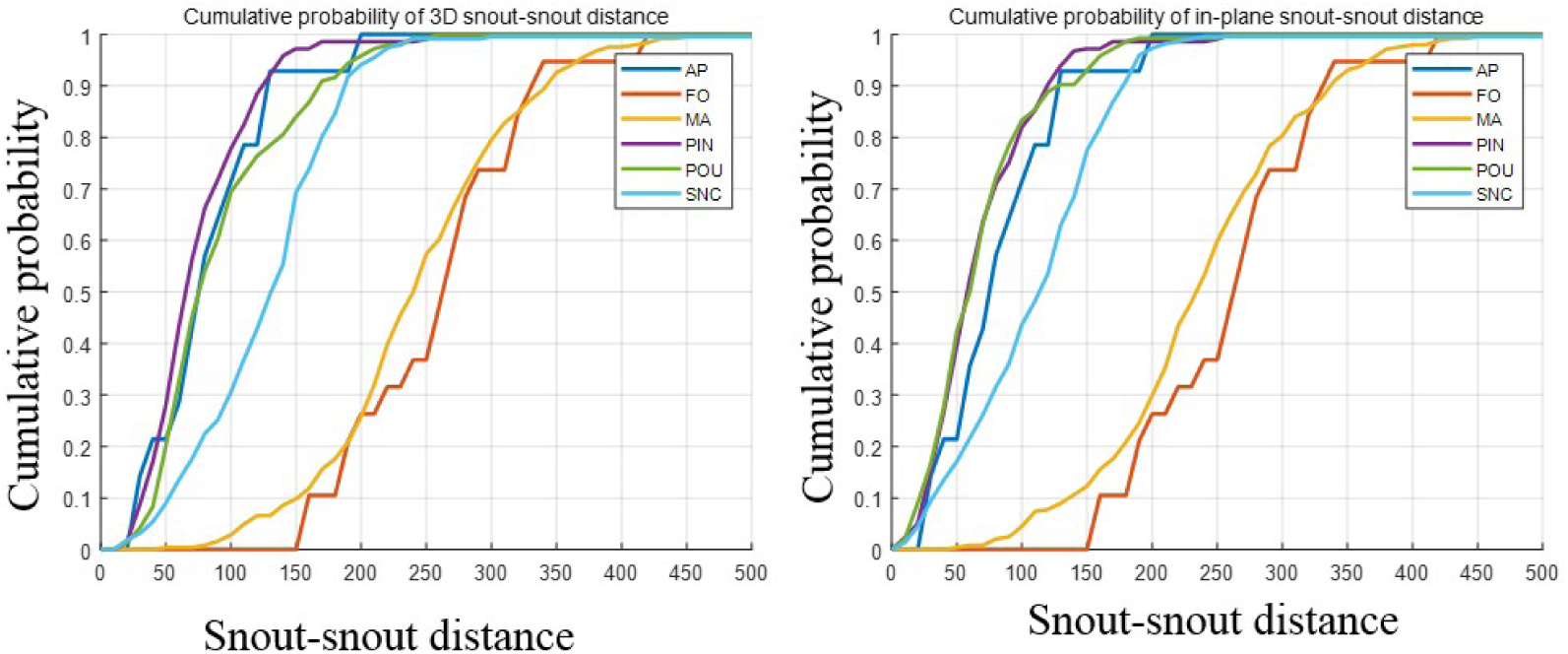
Quantitative evaluation of the snout-snout distance distribution for rat-rat free interaction scenarios. For social behaviors (PIN, POU, SNC), the cumulative probability of rat 3D and in-plane snout-snout distance less than 50 mm and 100 mm are (28.1%, 20.0%, 9.0%), and (39.6%, 42.3% and 17%) respectively.

**Figure 3—figure supplement 5.**
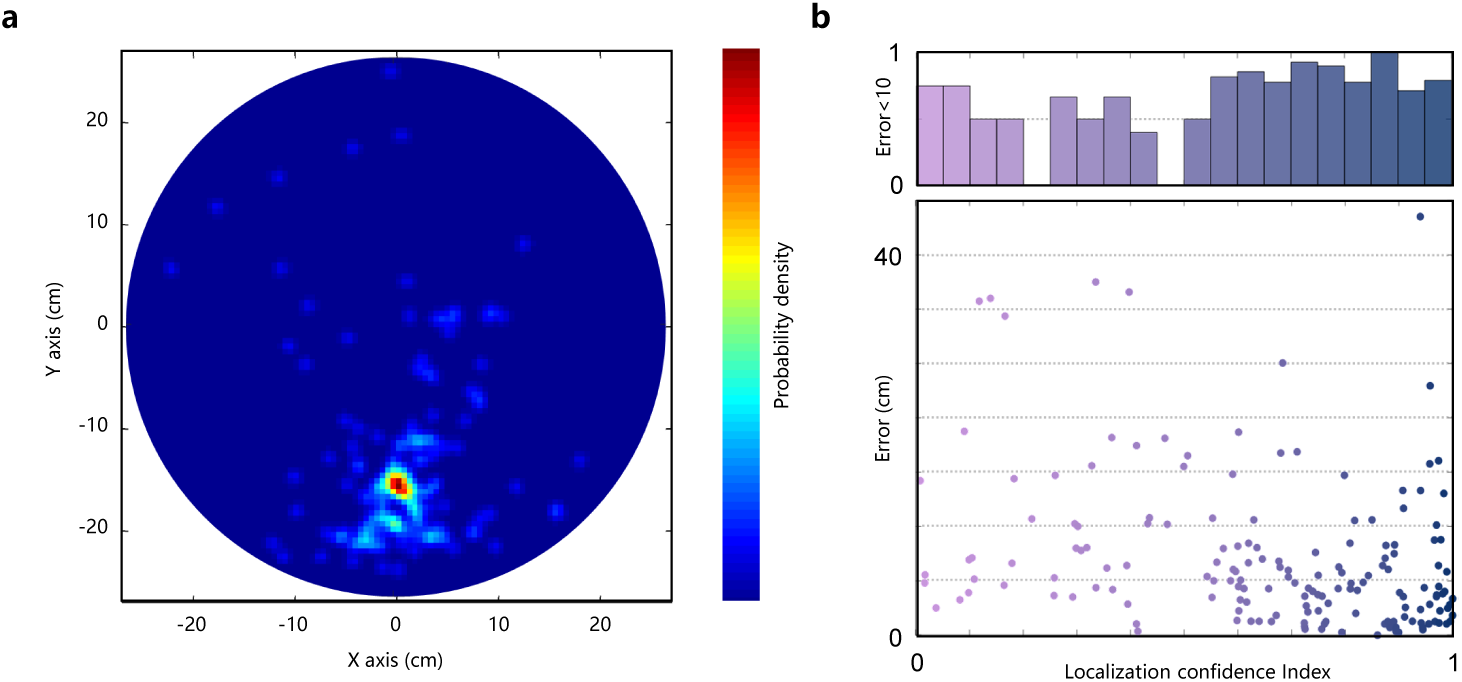
Validating ARBUR:SSL. **a**, The probability density map of the results estimated by ARBUR:SSL. USVs of the rat during still were used for evaluation. The ground truth of the horizontal coordinate of the rat nose is (0.5, -14.5), near the maximum of the density map. b, the SSL error (bottom), and the percentage of the results with an error less than 10 cm (top) versus the localization confidence index (LCI).

**Figure 4—figure supplement 1.**
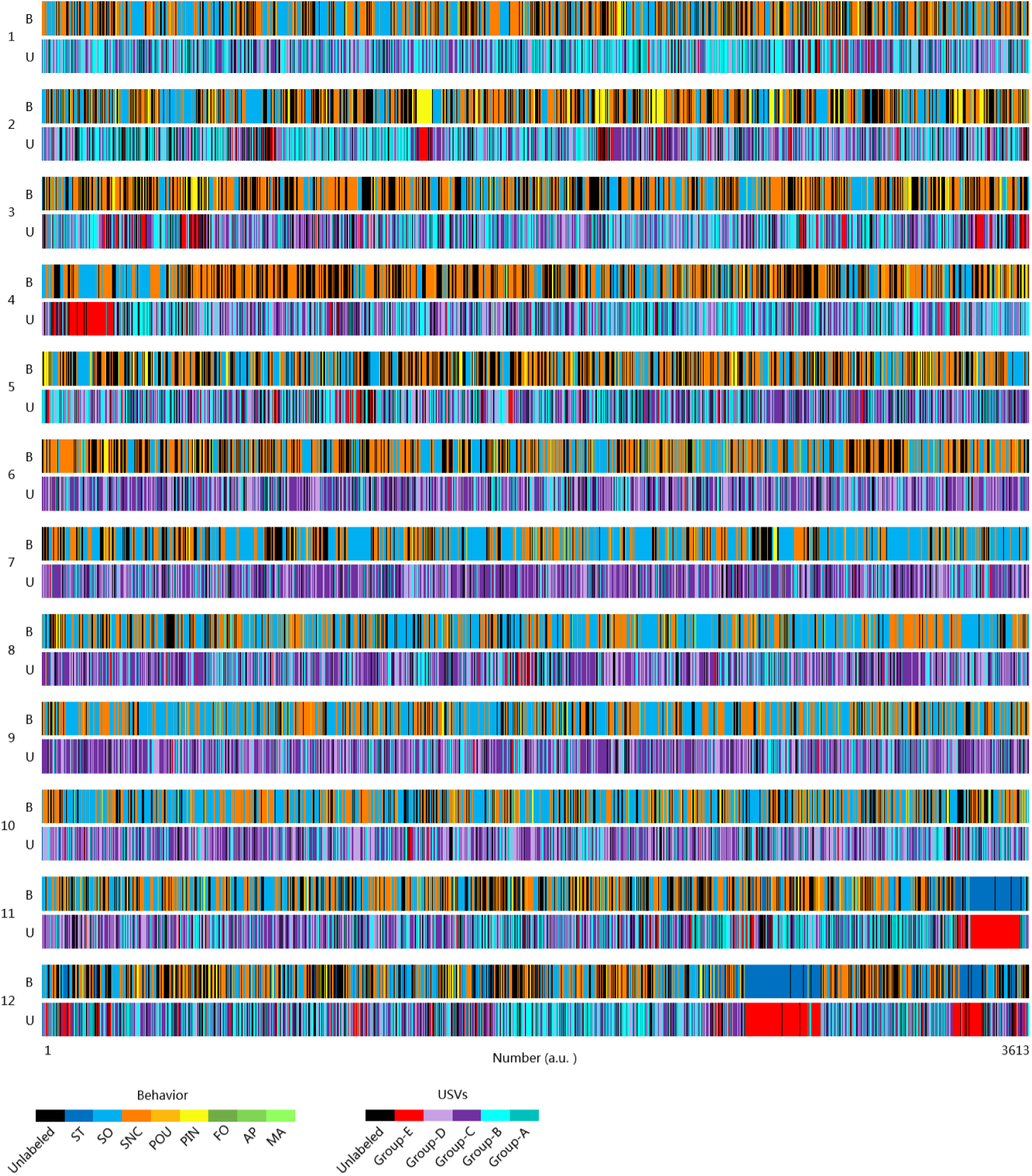
The behavior (B, top panels) and USV (U, bottom panels) ethograms of the simultaneously recorded vocal and behavioral repertoires. ST: still; SO: solitary; MA: moving away; AP: approaching; FO: following; PIN: pinning; POU: pouncing; SNC: social nose contact. Groups A and B: high-frequency non-step/step appetitive 50-kHz USVs; Groups C and D: low-frequency non-step/step appetitive 50-kHz USVs; Group E: aversive 22-kHz USVs.

